# Immunodominant structural proteins Gc and N drive T cell-mediated protection against La Crosse virus

**DOI:** 10.1101/2025.02.25.640063

**Authors:** Reem Alatrash, Bobby Brooke Herrera

**Affiliations:** Rutgers Global Health Institute, Rutgers University, New Brunswick, NJ, USA; Department of Medicine, Division of Allergy, Immunology, and Infectious Diseases and Child Health Institute of New Jersey, Rutgers Robert Wood Johnson Medical School, New Brunswick, NJ, USA

## Abstract

La Crosse virus (LACV) is a major cause of pediatric encephalitis in the U.S., primarily affecting children under 16 years. Despite severe morbidity and potential mortality risks, no vaccines or antivirals exist. Murine models recapitulate human susceptibility patterns, with weanling mice (3 weeks old) succumbing to disease while adults (≥8 weeks old) exhibit resistance. We characterized the T cell responses underlying this difference. Adult mice mounted robust CD4^+^ and CD8^+^ T cell responses by 6 days post-infection (dpi), with sustained IFN-γ, granzyme B, IL-2, and TNF-α production. These T cells expanded significantly and exhibited in vivo cytotoxicity against cells pulsed with LACV glycoprotein (Gc) and nucleocapsid (N) antigens. In contrast, weanlings showed weak T cell responses and 100% mortality by 7 dpi. Immunization with LFn-LACV-Gc and -N improved T cell cytotoxicity and survival in weanlings, highlighting their potential as vaccines. These findings inform strategies to mitigate LACV-induced encephalitis in children.

## INTRODUCTION

La Crosse virus (LACV) is a single-stranded, negative-sense RNA virus belonging to the *Bunyavirales* order ^1^. Its genome is segmented into three strands referred to as small (S), medium (M), and large (L), distinguished by their relative sizes. The S segment encodes the nucleocapsid protein (N) and nonstructural protein s (NSs), the latter of which has been shown to modulate the host cell antiviral response through innate immune pathways ^2^. The M segment encodes a glycosylated polyprotein precursor (GPC) that is cleaved by host cell proteases to generate the envelope glycoproteins Gn and Gc ^3^. Cleavage of GPC also produces the nonstructural protein m (NSm), which has been implicated in viral assembly ^2^. The L segment encodes an RNA-dependent RNA polymerase (RdRp), responsible for both transcription and replication of the viral genome ^4^.

Human infection with LACV occurs predominantly through the bite of infected Eastern tree hole mosquitoes (*Aedes triseriatus*), which are endemic throughout North America. Other mosquito species can also harbor LACV, potentially facilitating viral spread into new regions and hosts ^5^. Following entry into the human host, LACV replicates in skin muscle cells, enabling subsequent viral dissemination to multiple tissues and organs ^6,7^. Although most LACV-infected individuals experience only mild, febrile symptoms, the virus can invade the central nervous system (CNS) causing a subset to experience severe neurological manifestations, including altered behavior, seizures, coma, or even death ^8^. Pediatric populations, particularly children under the age of 16 years, exhibit the highest susceptibility to LACV encephalitis, implicating age-associated factors in disease severity ^9^.

Currently, there are no approved antiviral therapies or vaccines against LACV, necessitating reliance on supportive care alone to mitigate disease symptoms. The absence of licensed vaccines partly reflects a limited understanding of how the host immune response mediates protection, particularly against neurological sequelae. This gap in knowledge has been difficult to address in humans, where clinical samples, especially those collected during the acute phase of infection, are challenging to obtain. However, murine models accurately recapitulate the age-dependent susceptibility seen in humans, with weanling mice (3 weeks old) succumbing to LACV-induced neuropathology, whereas adults (≥8 weeks old) resist peripheral LACV infection ^6,10,11^.

Previous studies in adult mice indicate a critical role of the peripheral immune system in preventing LACV neuroinvasion, given that inoculation of C57BL/6 wild type mice via intraperitoneal (IP) or intradermal (ID) routes does not induce overt neurological disease, whereas intracranial inoculation does ^7,12-14^. Additional research reveals that type I interferon responses are indispensable for protecting mice from lethal peripheral LACV challenge at all ages, highlighting the importance of innate immunity in early antiviral control ^13,15^. Furthermore, in peripheral tissues, activation of antiviral recognition pathways induces type I interferon responses, such as IFN-β, which suppress viral replication, control LACV infection, and limit its spread to the CNS ^15,17,18^. However, within the CNS, similar signaling can also lead to neuronal damage and cell death ^19^. This dual role of innate immune responses in LACV infection underscores their context-dependent effects, providing both protection and potential pathogenicity ^16^.

While innate immune responses to LACV have been well studied, much less is known about adaptive immunity. Research has shown that myeloid dendritic cells, which bridge innate and adaptive immunity, are critical for early protective responses in adult mice infected with LACV ^13^. Their role is further enhanced in young mice upon treatment with type I interferon ^13^. Notably, both innate and adaptive immune responses appear to cooperate in controlling LACV infection in adult mice. The depletion of CD4^+^ and CD8^+^ T cells in adult mice reverse their resistance phenotype, emphasizing that T cell-mediated responses are critical for protection and prevention of neurological complications ^12^. This study also highlighted that while lymphocytes, including NK cells and CD4^+^ and CD8^+^ T cells infiltrate the CNS of susceptible weanling mice, their depletion does not impact neurological disease, suggesting they are not contributors to pathogenesis in young mice ^20^. Conversely, in adult mice, the depletion of both T cells and B cells results in increased viral loads in the brain and heightened neurological disease ^20^. Despite this work, the precise immunological correlates of protective immunity remain insufficiently characterized, particularly with respect to identifying virus-specific T cell responses and their antigenic targets.

We recently demonstrated that LACV-infected weanling mice produce a Th2-skewed cytokine profile and exhibit few IFN-γ producing T cells as compared to adults ^21^. However, the passive transfer of splenic leukocytes from immune adult mice to naïve weanlings prior to LACV challenge significantly improved survival ^21^. We also recently showed that adult mice have delayed spread of LACV into the CNS as compared to weanlings, further supporting adaptive immunity in peripheral control of the virus ^22^. We hypothesize that T cell-mediated immunity is essential for protection against LACV neuropathogenesis. Thus, a better understanding of age-dependent T cell responses is pivotal for designing effective vaccines and immunotherapies that target pediatric populations.

In this study, we provide a detailed analysis of T cell responses in adult and weanling mice across the LACV proteome. We demonstrate robust, polyfunctional CD4^+^ and CD8^+^ T cell responses directed against both structural and non-structural proteins in LACV-infected adult mice, whereas these responses are significantly blunted in weanlings. We additionally show that upon immunization structural proteins Gc and N elicit high in vivo cytotoxic activity in weanling mice that significantly improve survival. These findings underscore the need for vaccine development targeting these immunodominant antigens and highlight key immune determinants that could be leveraged to protect children from LACV-induced encephalitis.

## RESULTS

### Adult mice mount robust LACV-specific cellular responses, while weanlings lack these responses

Previous studies have documented high immunogenicity of structural antigens in bunyavirus infections, but the immunogenicity of non-structural proteins has been less thoroughly investigated ^1^. To identify LACV T cell targets, we performed in silico peptide analysis of the entire LACV proteome, evaluating potential epitopes restricted by both human and mouse MHC Class I and Class II molecules. Our analysis highlighted multiple immunogenic peptides in structural and non-structural proteins. Out of 176 total peptides identified as binders to MHC Class I molecules, an average of 12, 18, and 7 peptides were associated with Gn, Gc, and N, respectively (Fig. S1). For the non-structural proteins, we identified 5, 6, and 59 peptides binding to NSs, NSm, and RdRp, respectively, suggesting that these proteins may also serve as dominant targets for T cell-mediated immunity (Fig. S1). Among the peptides identified, Gc and RdRp contained the highest number of highly immunogenic sequences. Notably, 12 peptides were predicted to bind to both human and mouse MHC Class I molecules, underscoring their potential as cross-species immunogenic epitopes (Table 1). Based on these predictions, we proceeded with ex vivo analyses using a panel of immunogens, consisting of a modified *Bacillus anthracis* lethal factor (LFn) fused to each of the LACV structural or non-structural proteins. LFn has been shown to transport full-length antigens into the cytosol to induce virus-specific T cell responses via the MHC Class I and Class II pathways ^23-26^.

**Table 1.**
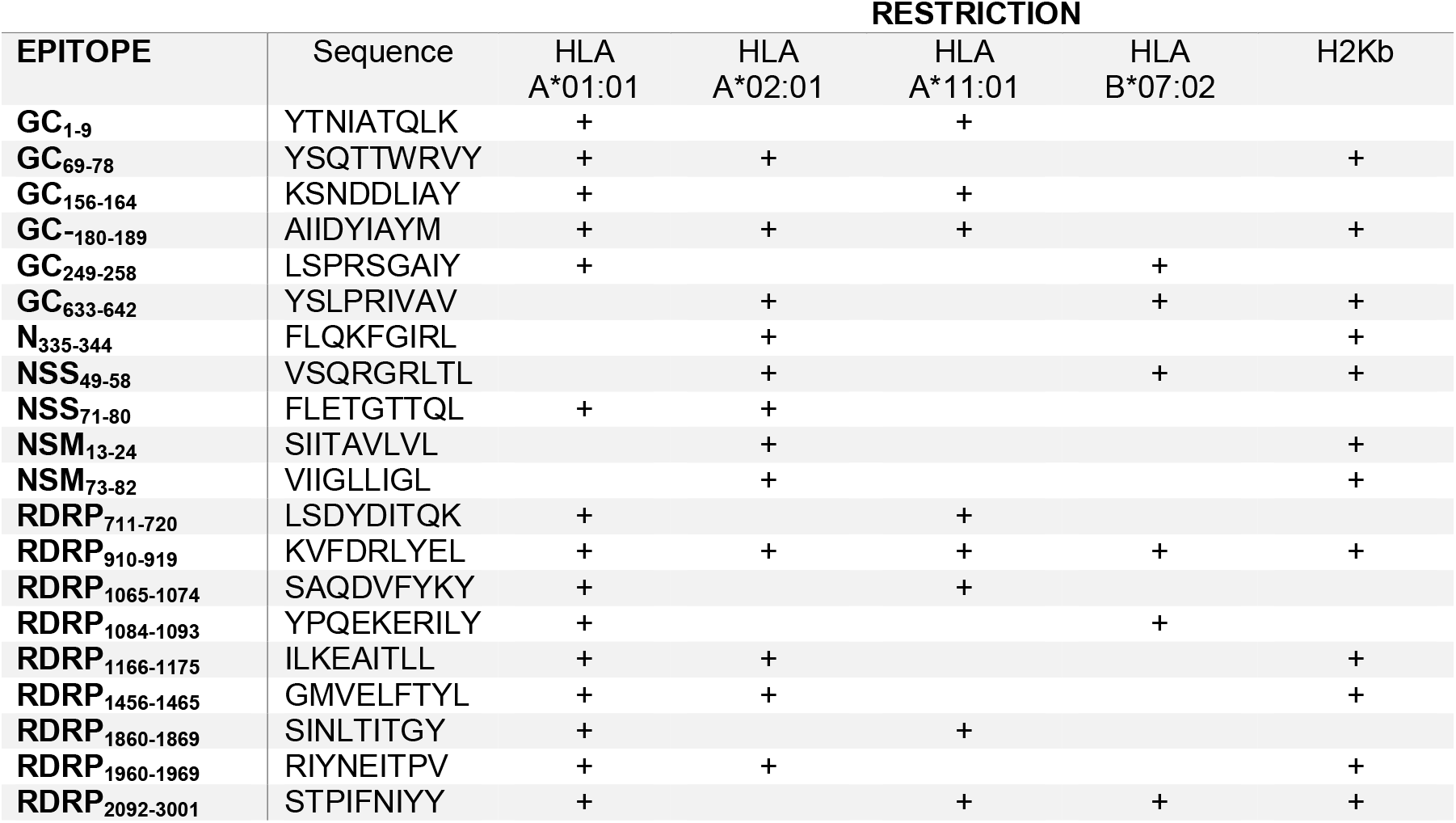
Predicted immunogenic LACV derived epitopes with human and/or MHC restriction.

To quantify the LACV-specific cellular responses, we measured IFN-γ production by splenic leukocytes from LACV-infected and uninfected C57BL/6 wild type mice in enzyme-linked immunospot (ELISPOT) assays at 6 and 14 days post-infection. Adult mice mounted a robust cellular response to multiple LACV proteins by 6 dpi, which persisted by 14 dpi (Fig.1A, S2A-B). Structural proteins Gc and N induced the strongest responses, followed by NSm as compared to the negligible responses observed in uninfected adult controls (Fig. 1B, S2A-B). In contrast, weanling mice at 6 dpi showed minimal IFN-γ responses to any LACV proteins, which were similar to the uninfected controls (Fig. 1A, S3A). Because all weanling mice succumbed to infection by 7 dpi, data at 14 dpi could not be obtained. These data indicate that the age-dependent immunological disparity in LACV infection extends to both the magnitude and breadth of T cell responses.

**Figure 1.**
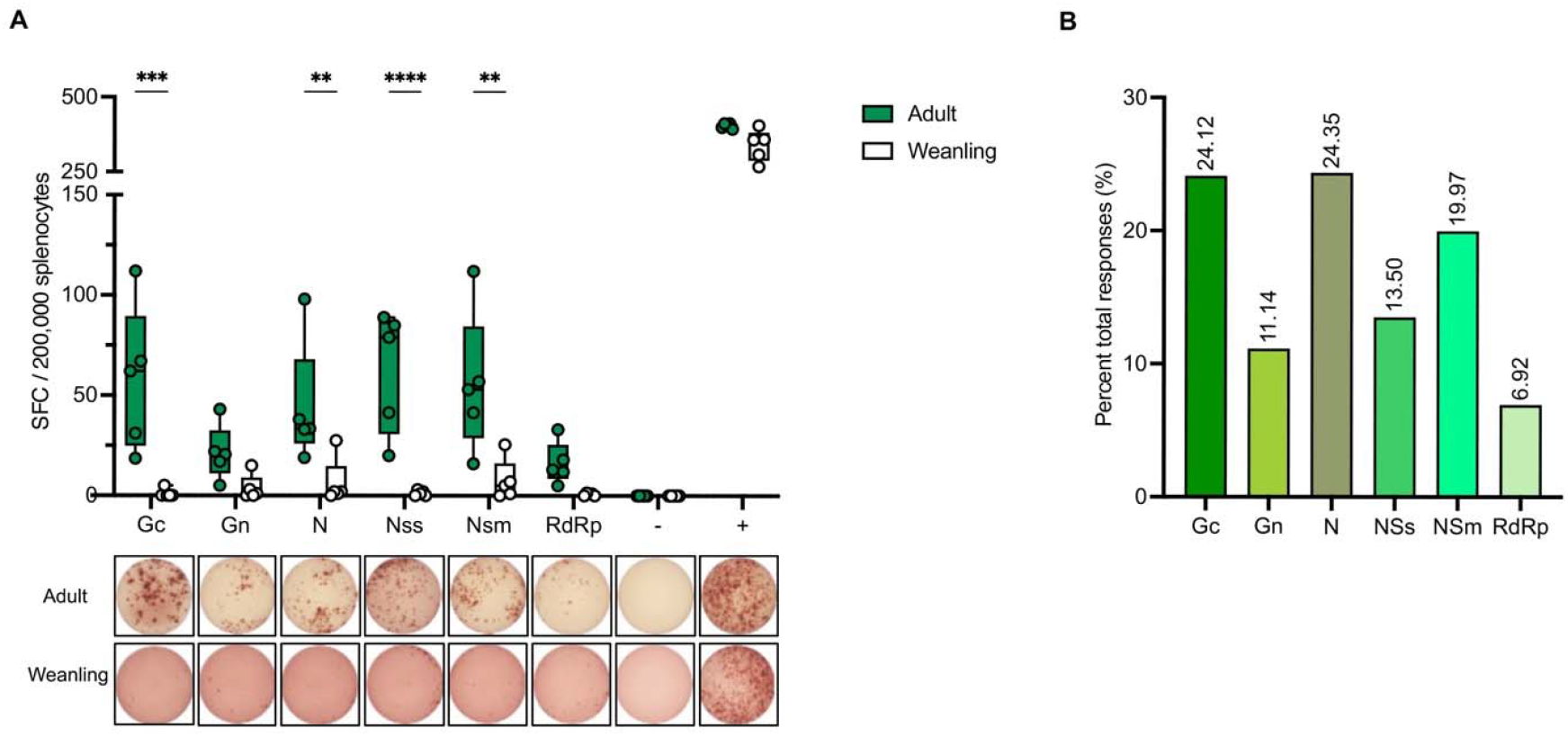
LACV-specific ex vivo cellular response in wildtype mice. (A) Top: Quantified spot forming cells (SFC) detected by ELISPOT assay using splenic leukocytes from adult and weanling mice infected with LACV at 6 days post-infection (dpi). Bottom: Splenic leukocytes were treated with LFn-LACV proteins and IFN-γ responses were detected by ELISPOT assay. (-) negative control: cells stimulated with LFn alone, (+) positive control: cells stimulated with PMA (phorbol 12-myristate 13-acetate). Statistical significance was determined using ANOVA test, *p < 0.05, **p < 0.01, ***p < 0.001, ****p < 0.0001. Data is presented as individual points with minimum and maximum values, (n=5 in each group). (B) Percent of total responses generated against each LFn-LACV protein in adult mice at 14 dpi. See also Figure S2, S3.

### Adult mice exhibit significantly higher percentage of both CD4^+^ and CD8^+^ splenic T cells following LACV infection compared to weanlings

We next evaluated whether the limited LACV-specific cellular responses in weanlings reflect a smaller pool of available T cells. Splenocytes from adult and weanling mice were analyzed by flow cytometry at 6 and 14 dpi for the frequency of CD4^+^ and CD8^+^ T cells. In adult mice, LACV infection triggered a significant increase in CD4^+^ T cell numbers at 6 dpi compared to age-matched uninfected controls at the same timepoint (Fig. 2A). This population expansion persisted through 14 dpi. Similarly, CD8^+^ T cells in adults exhibited a more pronounced expansion, reaching statistical significance at both time points (Fig. 2B). By contrast, the percentage of CD4^+^ and CD8^+^ T cells in weanlings at 6 dpi remained unchanged from uninfected age-matched controls and significantly lower percentages observed in infected adult mice at 6 dpi, corresponding with their poor ex vivo IFN-γ response. These findings suggest that both CD4^+^ and CD8^+^ T cell numbers expand in adults but not weanlings post LACV infection.

**Figure 2.**
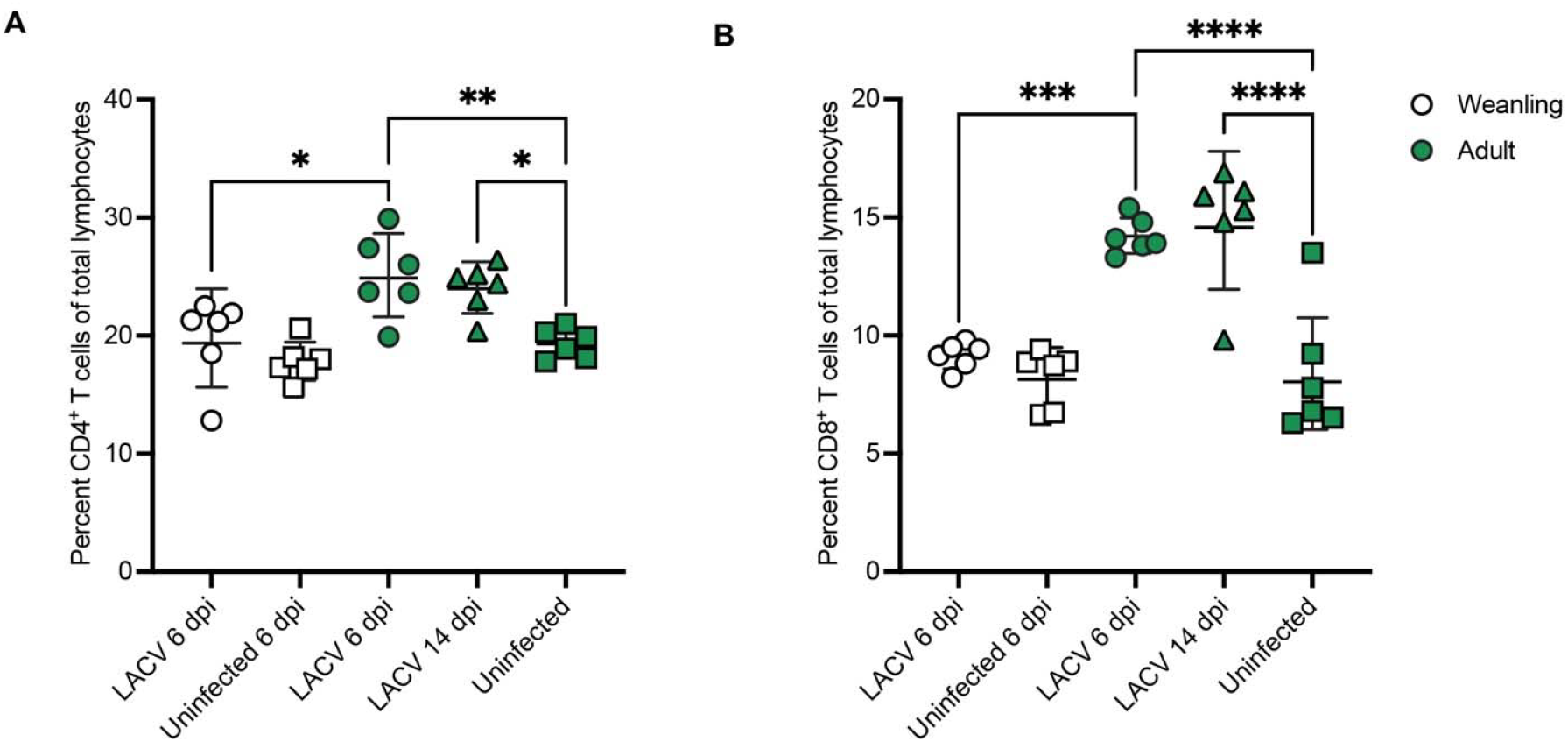
LACV-induced CD4^+^ and CD8^+^ T cell Expansion in LACV infected mice. Total percentage of CD4^+^ T cells in (A) and CD8+ T cell populations (B) among total splenocytes of adult and weanling mice at 6 and 14 days post-infection (dpi) with LACV analyzed by flow cytometry. Statistical significance was determined using ANOVA test, *p < 0.05, **p < 0.01, ***p < 0.001, ****p < 0.0001. Data is presented as individual points with mean ± standard deviation (n=6 in each group).

### Adult mice mount functional LACV-specific CD4^+^ and CD8^+^ T cell responses, whereas weanlings do not

To determine the functional capacity of virus-specific T cells in adult and weanling mice following LACV challenge, we performed intracellular cytokine staining of splenic leukocytes pulsed with LFn-LACV immunogens. We simultaneously measured the production of IFN-γ, granzyme B, IL-2, and TNF-α in CD4^+^ and CD8^+^ T cells (Fig. 3, S4-8). Adult mice displayed robust IFN-γ responses across structural and most non-structural proteins by 6 dpi. In adult mice, CD4^+^ and CD8^+^ T cells displayed a robust IFN-γ response to all structural and most non-structural LACV proteins as early as 6 dpi, with levels significantly higher than in age-matched uninfected controls (Fig. 3A-B). CD4^+^ T cell responses began to decline by 14 dpi, while CD8^+^ T cell responses increased at 14 dpi against most LACV proteins, indicating sustained effector activity of CD8^+^ T cells (Fig. 3A-B). Importantly, weanling mice exhibited negligible cytokine responses by CD4^+^ and CD8^+^ T cells, mirroring their poor splenic T cell expansion and heightened mortality (Fig. 3C-D, S7).

**Figure 3.**
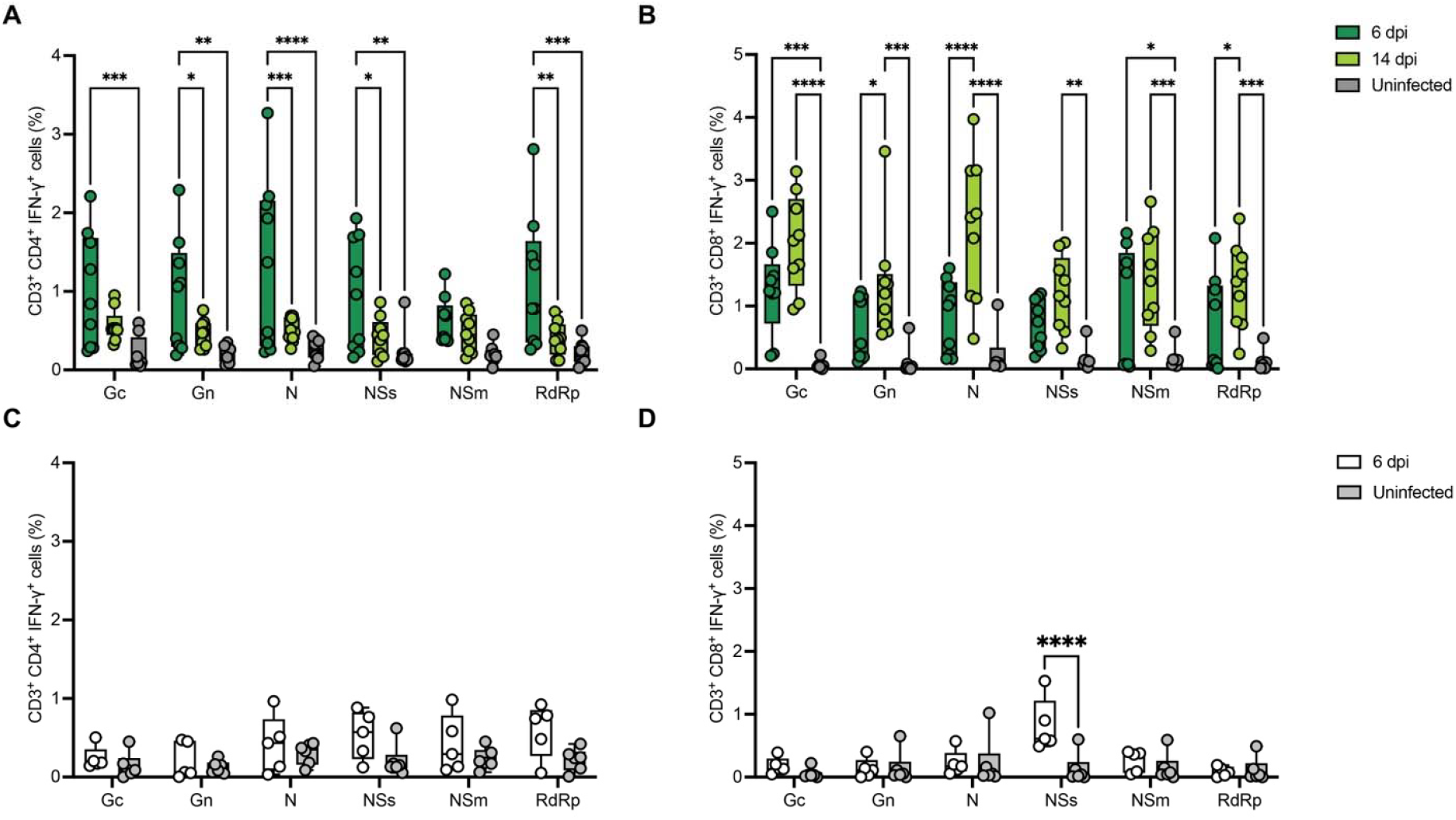
LACV-specific ex vivo splenic CD4^+^ and CD8^+^ T cell Functional response in LACV infected mice. Functional LACV-specific CD4^+^ and CD8^+^ T cell responses shown as the percent of CD4^+^ or CD8^+^ cells positive for IFN-γ in CD3^+^ gate in flow cytometry/intracellular cytokine staining in adult (A, B) and weanling mice (C, D) at 6, 14 dpi compared to age-matched uninfected controls. Statistical significance was determined using ANOVA test, *p < 0.05, **p < 0.01, ***p < 0.001, ****p < 0.0001. Data is presented as individual points with minimum and maximum values, (n=9 adults, 5 weanlings). See also Figure S4-8.

A similar trend was observed for other cytokines. Granzyme B expression was significantly elevated in adult mice but was minimal in weanlings (Fig. S4, S7). IL-2 and TNF-α responses were generally weak across both age groups, though adult mice consistently showed higher levels than weanlings (Fig. S5-7). TNF-α expression was detectable in adults and weanlings but remained low overall compared to other analyzed cytokines (Fig. S6). These results indicate that virus-specific T cells in adult mice mount functional effector responses during a LACV infection, whereas such responses are notably weak or absent in weanling mice.

We further examined polyfunctionality by assessing the capacity of T cells to concurrently produce multiple cytokines. In adult mice, CD4^+^ T cells exhibited multi-cytokine functionality, although IFN-γ was the dominant effector molecule (Fig. 4A-B). These CD4^+^ T cell effector functions were sustained at 14 dpi (Fig. S9A-B), underscoring their role in orchestrating immune responses during LACV infection. Furthermore,∼20–34% of CD8^+^ T cells co-produced IFN-γ and granzyme B at 6 dpi, rising to 54-69% by 14 dpi. (Fig. 4C-D). A smaller proportion of CD8^+^ T cells demonstrated dual functionality involving granzyme B with IL-2 or IFN-γ with IL-2 (Fig. 4C-D). These polyfunctional responses were most pronounced following pulsing cells with the LFn-LACV Gc, Gn, or RdRp immunogens, underscoring the immunogenicity of these proteins (Fig. 4A-B, S9). Thus, in contrast to weanlings, adult mice generated a robust, polyfunctional T cell response that likely contributes to effective viral clearance and survival.

**Figure 4.**
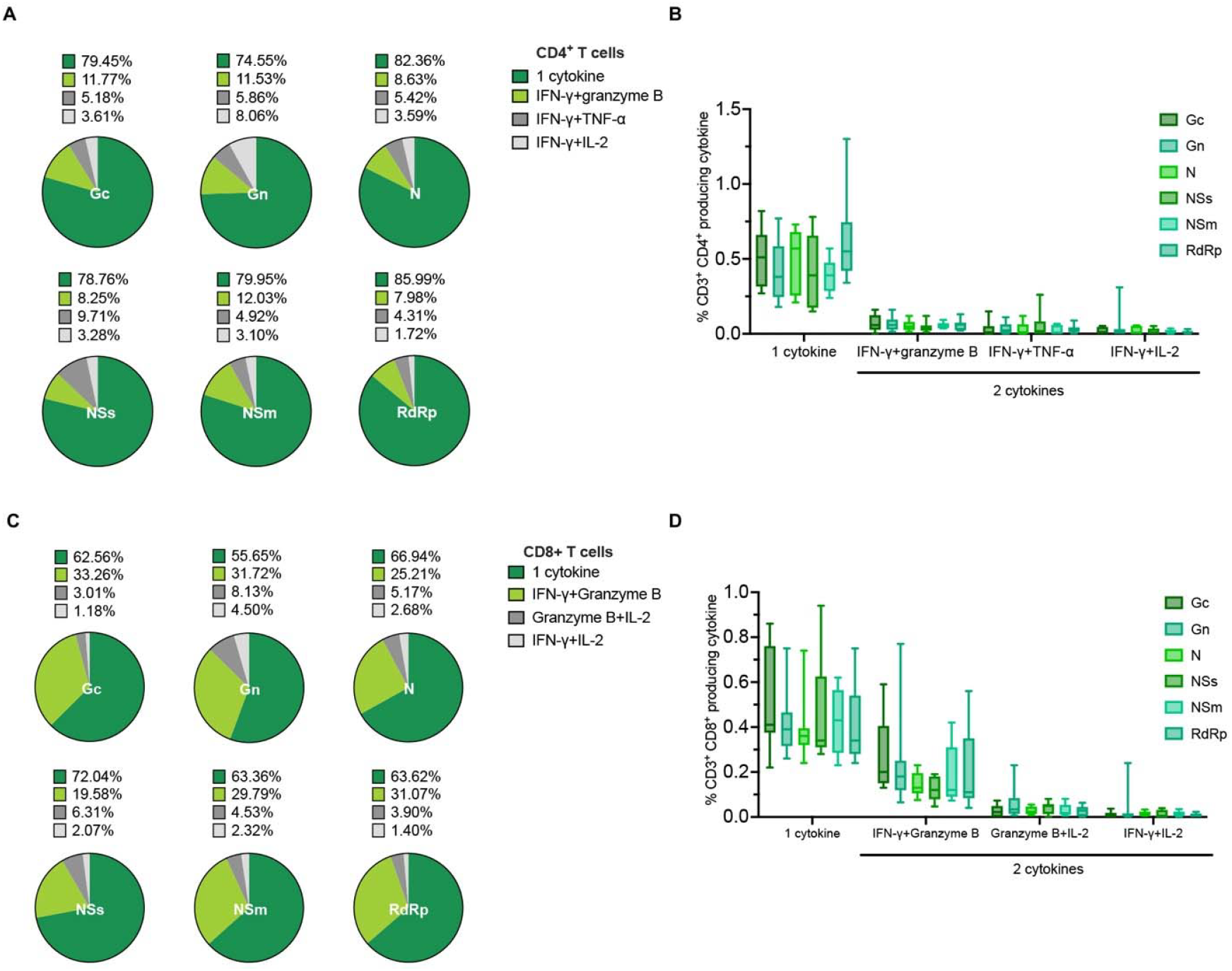
Polyfunctional profiles of LACV-specific CD4^+^ and CD8^+^ T cells at 6 dpi. Pie charts depicting the distribution of LACV-specific CD4^+^ T cell (A) and CD8^+^ T cell (C) populations with single or dual functional profiles, defined by cytokine production (n=9). (B, D) Bar graph showing the total percentage of LACV-specific CD4^+^ T or CD8+ T cells producing one or two cytokines in response to infection. Data is represented as mean with minimum and maximum values (n=9). See also Figure S9.

### T cell responses against LACV Gc and N exhibit potent in vivo cytotoxicity and confer protection in weanlings upon vaccination

We next assessed T cell cytotoxic activity using an in vivo cytotoxicity assay based on carboxyfluorescein succinimidyl ester (CFSE). Briefly, CFSE-labeled splenocytes pulsed with individual LFn-LACV proteins (LFn-LACV-Gc or -N), were transferred into LACV-infected adult mice at 14 dpi, and LACV-infected weanlings at 6 dpi, and the percentage of target cell killing was calculated (Fig. 5A). In adult mice, target cells pulsed with LFn-LACV-Gc or -N were eliminated at rates of ∼53% and ∼40%, respectively (Fig. 5A, 5D). In contrast, weanling mice exhibited minimal killing of target cells pulsed with LFn-LACV-Gc (∼5%) or LFn-LACV-N (∼7%) at 6 dpi, the latest timepoint that could be assessed (Fig. 5A, D). These results show that LACV-specific T cells mediate cytotoxicity in LACV-infected adult mice but not in weanlings.

**Figure 5.**
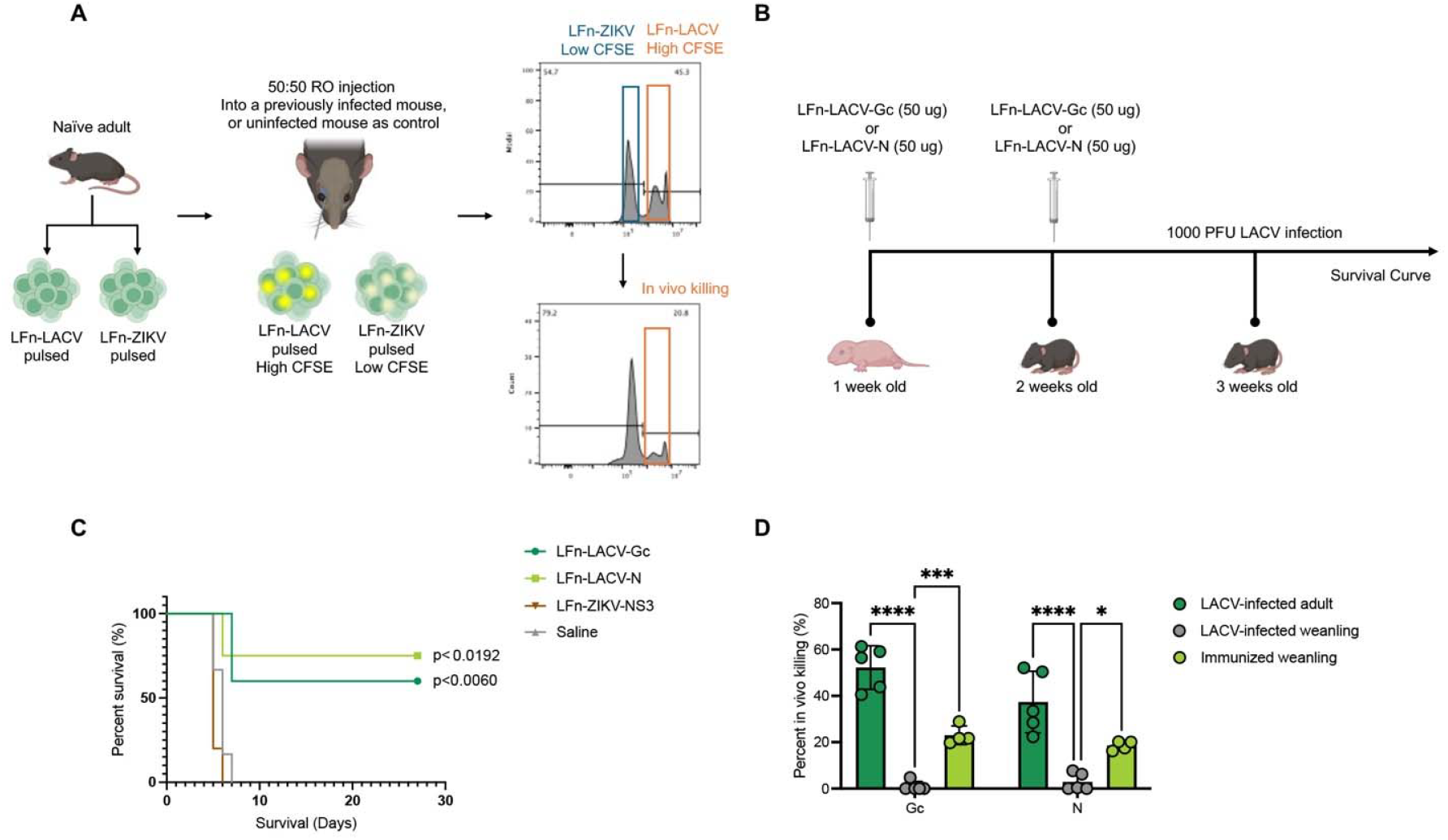
In vivo cytotoxicity and protection mediated by LACV-specific T cells. (A) Schematic representation of the in vivo cytotoxicity assay. Equal numbers of naïve splenocytes pulsed with LFn-LACV-Gc or LFn-LACV-N (CFSE^High^ target cells) and splenocytes pulsed with an irrelevant protein, LFn-ZIKV-NS3 (CFSE^Low^ marker cells), were transferred into either naïve or LACV-immune recipient mice. Percent killing was quantified by flow cytometry in recipient spleens 4 hours post-transfer. (B) Schematic of the LFn-LACV immunization strategy. One-week-old mice were immunized with 50 µg of either LFn-LACV-Gc or LFn-LACV-N. A booster dose was administered 7 days later, followed by LACV challenge at weanling age. (C) Survival curves of weanling mice immunized with LFn-LACV-Gc or LFn-LACV-N compared with LFn-ZIKV-NS3 and subsequently challenged with LACV (n = 5 per group). Statistical significance was determined using log-rank test. (D) Bar chart quantifying in vivo cytotoxicity in previously LACV-infected adult and weanling mice, as well as in LFn-LACV-Gc- or LFn-LACV-N-immunized weanling mice. Percent killing was determined by flow cytometry. Statistical significance was determined using ANOVA test, *p < 0.05, **p < 0.01, ***p < 0.001, ****p < 0.000. Data is presented as individual points with mean ± standard deviation (n=5 per group).

To further translate these findings into a vaccination strategy for susceptible weanling mice, we immunized neonatal mice (1 week old) with either LFn-LACV-Gc or -N, with a boosting dose at 2 weeks old, then subsequently challenged them with LACV at weanling age (Fig. 5B-C). The two-dose immunization regimen significantly improved survival compared to control groups, increasing survival rates to ∼70–80% (Fig. 5C). To assess whether immunization impacted T cell cytotoxicity in weanlings, we repeated the CFSE-based in vivo cytotoxicity assay post-immunization. CFSE-labeled splenocytes pulsed with individual LFn-LACV proteins were transferred into their respective LFn-LACV-Gc or -N immunized recipient mice, and cytotoxicity was measured. Strikingly, LFn-LACV-Gc immunization enhanced target cell killing to ∼21% in weanling mice, a significant increase compared to the negligible cytotoxic response observed post-infection. Similarly, LFn-LACV-N immunization improved target cell elimination to ∼19%, demonstrating a clear enhancement in cytotoxic capacity relative to unvaccinated weanling mice (Fig. 5D). These results confirm that vaccine-elicited T cell responses contribute to protection against LACV-induced disease in weanling mice by enhancing T cell cytotoxic activity.

## DISCUSSION

In this study, we characterized virus-specific T cells in adult and weanling mice providing a critical step toward developing a pediatric vaccine for this clinically significant arbovirus. Our findings demonstrate significant differences in T cell responses between adult and weanling mice. Adult mice mounted robust LACV-specific T cell responses, characterized by strong IFN-γ production against both structural and nonstructural antigens, with the highest responses targeting the structural proteins Gc and N. In contrast, weanling mice displayed significantly weaker T cell responses at 6 dpi, underscoring the inability of their immature immune systems to mount effective adaptive cellular immunity. This disparity likely contributes to the heightened susceptibility of weanlings to severe LACV infection.

Further analysis revealed significant expansion of CD4^+^ and CD8^+^ T cell populations in adult mice following LACV infection, accompanied by robust cytokine production, including IFN-γ, granzyme B, IL-2, and TNF-α. Polyfunctional CD4^+^ and CD8^+^ T cells, which co-produced cytokines especially IFN-γ and granzyme B, were prominent in adults, supporting effective viral control and clearance. By contrast, weanlings exhibited limited T cell expansion and functional responses, suggesting an immature adaptive immune system unable to mount sufficient antiviral defenses.

Consistent with prior studies, our findings reaffirm the essential role of CD4^+^ and CD8^+^ T cells in mediating protection in adult mice ^20^. Depletion or knockout of these subsets led to disease progression and neurological complications at later stages of LACV infection in adult mice, emphasizing the necessity of adaptive immunity in sustaining the protective effects initially provided by innate responses ^20^. On the other hand, the poor immune outcomes in weanlings may stem from deficiencies in both innate and adaptive immunity in younger mice. Evidence suggests that dendritic cells in young mice exhibit impaired functionality, potentially due to reduced maturity or lower expression of co-stimulatory molecules ^27^. Additionally, a limited T cell receptor (TCR) repertoire, immature germinal center reactions, and suboptimal T follicular helper (Tfh) cell activity further compromise the adaptive immune response in weanlings ^27^. Understanding these mechanisms is critical for informing vaccine strategies aimed at enhancing immunity in vulnerable populations.

In our efforts to inform vaccine design, we explored the immunogenicity against the entire LACV proteome. Our ex vivo analysis, which included T cell analysis against both structural and nonstructural proteins, represents a significant advancement over previous ex vivo studies on members of *Bunyavirales* which focused mostly on structural antigens ^1^. Immunogenic epitopes were identified across the proteome, providing a broad array of potential T cell targets. The strong immunogenicity of the structural proteins Gc and N was validated through their ability to elicit robust T cell responses with significant in vivo cytotoxicity. Moreover, immunization with these proteins conferred protection in weanling mice upon LACV challenge, demonstrating their potential as vaccine candidates. These findings align with studies on genetically related viruses within the *Peribunyaviridae* family, such as Jamestown Canyon Virus (JCV), Oropouche Virus (OROV), and Bunyamwera Virus (BUNV), where glycoproteins (Gn/Gc) and nucleocapsid (N) proteins were identified as key targets for T cell responses based solely on immunoinformatic analysis ^28-30^. Similarly, comprehensive in silico analyses of Rift Valley Fever Virus (RVFV) and Severe Fever with Thrombocytopenia Syndrome Virus (SFTSV) have highlighted the immunogenicity of structural and nonstructural proteins ^31,32^. Notably, cross-reactive N-specific CD8^+^ T cells have been observed across hantaviruses, and T cell responses targeting Gn and Gc have been identified in patients infected with Hantaan virus (HTNV) and Andes Virus (ANDV) ^33-35^.

While the current study did not fully determine the specific mechanisms of vaccine-induced protection, we have established a foundation for future research to identify correlates of immunity for LFn-LACV-Gc or - N vaccines. These findings served as a validation of the high immunogenicity of identified viral proteins in this study and their protective potential in weanling mice. Notably, this work represents one of the few attempts to develop a vaccine against LACV, building on early studies that demonstrated protection using DNA vaccines encoding Gc/Gn and N proteins in adult *IFNAR1*^*-/-*^ mice ^36^. Mice vaccinated with DNA encoding the glycoproteins Gn and Gc produced neutralizing antibodies and exhibited a high degree of protection against challenge with high doses of LACV ^36^. Depletion of CD4^+^ T cells in mice vaccinated with DNA encoding Gn/Gc reduced their capacity to control the infection ^36^.

While most prior studies have utilized mouse models to investigate LACV pathogenesis, our understanding of T cell responses in LACV-infected humans remains limited. Surveillance data from LACV-endemic regions reveal that a majority of individuals with detectable neutralizing antibodies do not report a history of clinically significant infection ^8^. Among those who do develop encephalitis, clinical manifestations range from mild aseptic meningitis to transient or permanent neurological complications ^37^. This variability in susceptibility to infection and disease severity suggests a potential role for MHC-restricted T cell responses. Supporting this hypothesis, an earlier study examining MHC antigen frequencies in patients with LACV encephalitis identified HLA-specific risk factors associated with disease, such as the prevalence of HLA-B49 and/or the absence of HLA-DR5 ^38^. These findings underscore the importance of T cell-mediated immunity in determining disease susceptibility and severity in human LACV infections. Although there is a convenient mouse model available for studying LACV, there is a critical need to understand human HLA restriction and disease susceptibility and to characterize HLA-specific T cell responses in both young and adult populations.

By emphasizing the age-dependent nature of T cell responses and identifying immunogenic viral targets, this work provides a foundation for the development of LACV vaccines and therapeutics. Future research should focus on addressing the mechanistic basis of weak T cell responses in juvenile models, including deficits in antigen presentation and T cell priming. Bridging these gaps will further inform vaccine strategies to mitigate the burden of LACV disease, particularly in children, the population most at risk.

### Limitations of the study

One limitation of this study is that the immunogenic peptides predicted through in silico analysis were not experimentally validated ex vivo or in vivo. While we identified the immunogenicity of full-length LACV proteins, which informed vaccine development, further validation of individual peptide immunogenicity could provide a more precise understanding of T cell responses, particularly in the context of human immunity. Future studies should focus on experimentally confirming these predicted epitopes to better assess their relevance for vaccine design and immune response profiling.

## RESOURCE AVAILABILITY

### Lead contact

Further information and requests for resources and reagents should be directed to and will be fulfilled by the lead contact, Bobby Brooke Herrera (BBH).

### Materials availability

This study did not generate new unique reagents.

### Data and code availability

Any additional information required to reanalyze the data reported in this paper is available from the lead contact upon request.

## ACKNOWLEDGMENTS

We would like to thank Rutgers Global Health Institute, Rutgers Robert Wood Johnson Medical School, and the Child Health Institute of New Jersey for their support. We would also like to thank the Foundation for Health Advancement for grant support (IFHA 17-24). The funders had no role in the design, data collection, data analysis, and reporting of this study.

## AUTHOR CONTRIBUTIONS

Conceptualization, R.A. and B.B.H.; formal analysis, R.A. and B.B.H.; investigation, R.A., S.I, V.V, J.D, B.B.H; resources, B.B.H.; data curation, R.A., S.I, V.V, J.D; writing—original draft preparation, R.A. and B.B.H.; writing—review and editing, R.A. and B.B.H.; visualization, R.A. and B.B.H.; supervision, B.B.H.; funding acquisition, B.B.H. All authors have read and agreed to the published version of the manuscript.

## DECLARATION OF GENERATIVE AI AND AI-ASSISTED TECHNOLOGIES

N/A

## SUPPLEMENTAL INFORMATION

**Document S1. Figures S1–S9**.

## STAR⍰METHODS

### KEY RESOURCES TABLE

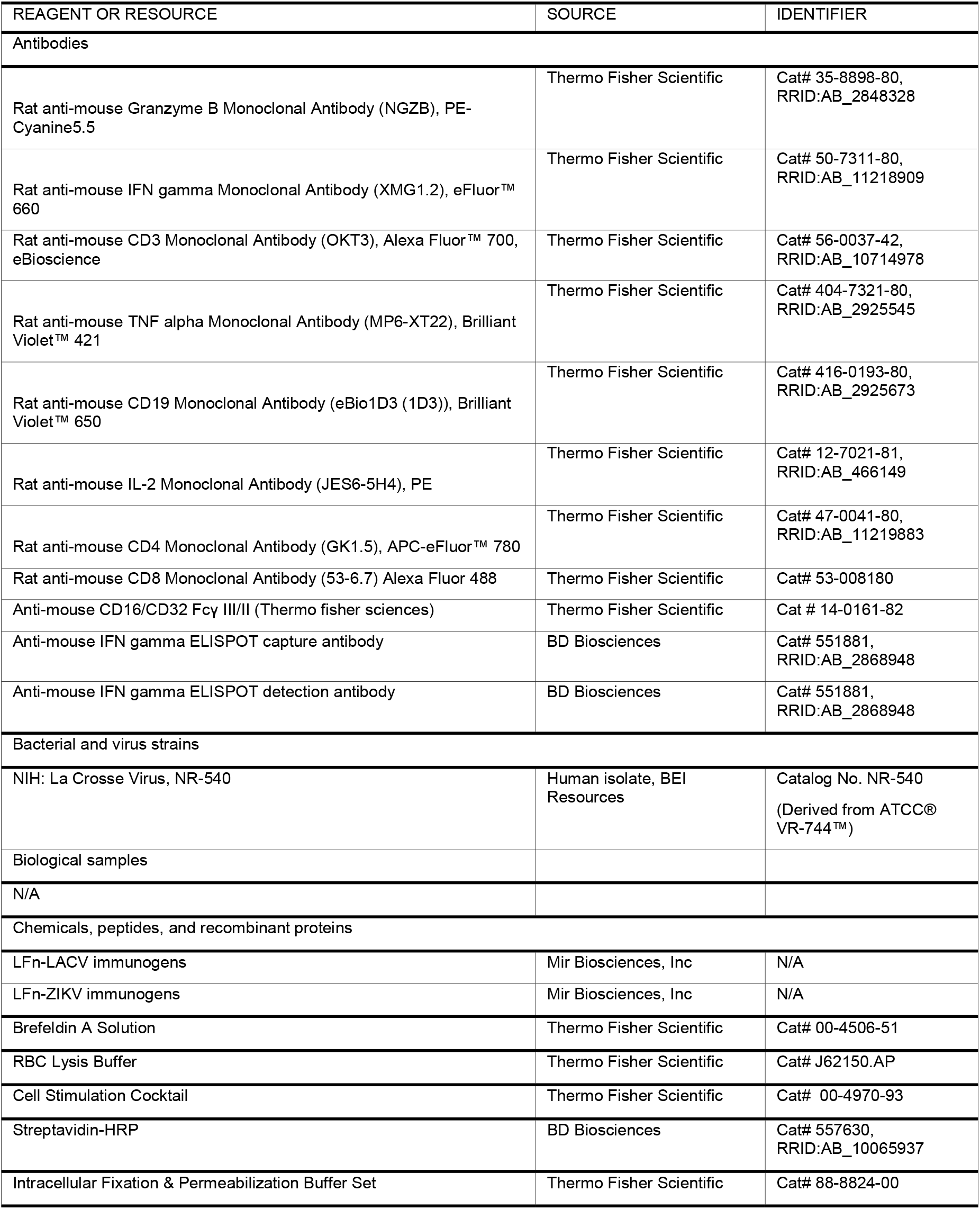
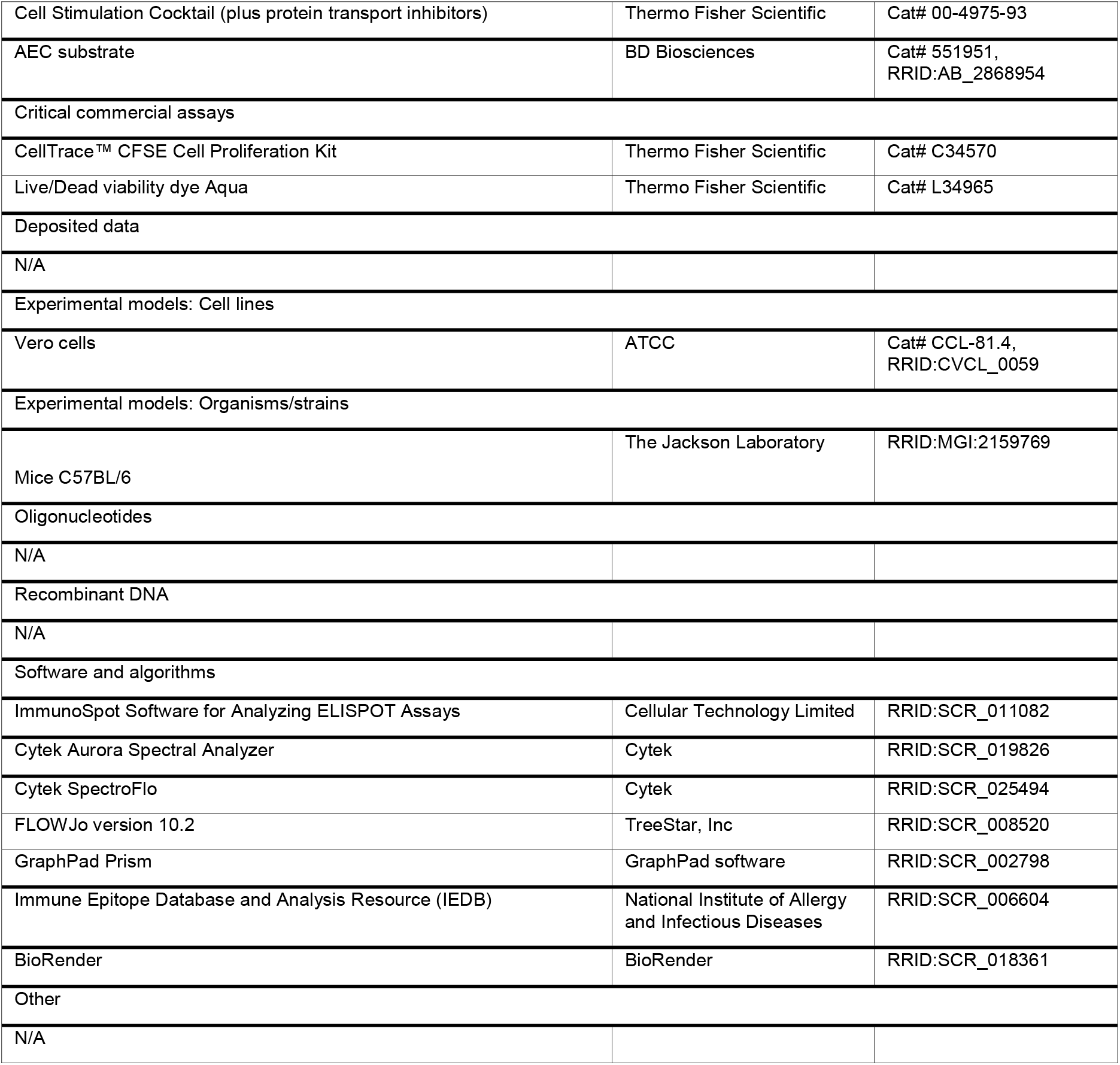

### EXPERIMENTAL MODEL AND STUDY PARTICIPANT DETAILS

#### Infection of mice with LACV and neurological disease progression criteria

All animal studies were approved by the Rutgers IACUC (protocol number 202300007). Wild-type (C57Bl/6) mice were purchased from Jackson Laboratories and maintained in a breeding colony at the Child Health Institute of New Jersey. LACV (La Crosse Virus, NR-540), a human isolate was purchased and obtained from BEI Resources. Mice at 3 (weanling) or 8 (adult) weeks of age were inoculated with 103 PFU of LACV in phosphate-buffered saline (PBS) intraperitoneally in a volume of 200 µL/mouse. Mice were observed daily for signs of neurological disease that included hunched posture, seizures, reluctance, or inability to move normally, or paralysis. Animals with clear clinical signs of neurological disease were scored as clinical and euthanized immediately.

### METHOD DETAILS

#### Bioinformatic analyses

All 9mer peptides encoded in the LACV proteome were predicted for their binding affinity to MHC class I alleles A*0201, A*0101, A*1101, and B*0702. C57BL/6 MHC I was also included for analyses (H2Kb). Binding predictions are performed using NetMHCpan 4.1 EL MHC class I prediction tool available on the Immune Epitope Database (IEDB) (http://www.iedb.org) Web site ^39^. Peptides are selected if they were in the top 1% of binders for each MHC molecule analyzed for further analysis for MHC I immunogenicity available on the IEDB website. Immunogenic peptides were predicted as previously described ^40^. Immunogenic peptides were assigned a positive immunogenic score. A total of 176 9-mer peptides were identified by MHC class I predictions, as described above. MHC II immunogenicity score of peptides predicted to bind using 7-allele method.

#### LFn-LACV immunogens

LFn-LACV immunogens were purchased from Mir Biosciences, Inc., (New Jersey, USA). In brief, gene fragments encoding the LACV Gc, Gn, N, NSm, NSs and RdRp were cloned into the LFn expression vector. The constructs were transformed into Escherichia coli BLR (DE3) (Millipore, Medford, MA). Selected transformants were sequenced to verify in-frame cloning. The LFn-LACV immunogens were expressed upon induction with isopropyl-β-d-thiogalactopyranoside (IPTG; Sigma-Aldrich, St. Louis, MO, USA) in 5 liters of Luria broth containing Kanamycin for 4 h. Cells were pelleted by centrifugation and resuspended in imidazole (1 mM) binding buffer (Novagen, Madison, WI) in the presence of a protease inhibitor cocktail (Thermo Fisher Scientific, Waltham, MA). Cell pellets were sonicated and centrifuged at 4°C, and the supernatants were loaded in an equilibrated nickel-charged column for affinity purification. The bound proteins were eluted in 100 to 200 mM imidazole, desalted with a Sephadex G-25M column (Sigma-Aldrich, St. Louis, MO, USA), and eluted in phosphate-buffered saline (PBS) (Sigma-Aldrich, St. Louis, MO, USA). The PBS-eluted proteins were passed through Detoxi-Gel (Thermo Fisher Scientific, Rockford, IL). Protein concentrations were determined, and samples were stored at −80°C.

#### Splenic leukocytes harvest

Spleens from LACV-infected and uninfected mice were harvested at 6 and 14 dpi, as described previously ^21^ .Spleens were processed into single-cell suspensions by passing them through a 70 µM mesh filter, followed by red blood cell lysis using RBC Lysis Buffer (Thermo Fisher Scientific). Isolated cells were then seeded for subsequent ELISPOT and flow cytometry/ICS assays.

#### ELISPOT test

Ex vivo ELISPOT tests ELISPOT assays were performed as previously described ^24^. Briefly, 96-well plates (Millipore) are coated with anti-mouse IFN-γ antibodies (BD Biosciences) and incubated overnight at 4°C. Plates are blocked with RPMI 10% FBS and washed 2 times. Splenic leukocytes harvested from LACV infected mice, as well as uninfected controls are seeded in triplicate at 2 x 105 cells/well. LFn-LACV immunogens are added to each well (5 μg/ml) and plates are incubated at 37°C with 5% CO2 for 18-24 hours. Cells stimulated with PMA (Phorbol 12-myristate 13-acetate) will serve as a positive control and cells stimulated with LFn alone as a negative control. After incubation and removal of cells, the detection antibody (BD Biosciences) is added and incubated at room temperature. Plates are washed then incubated with streptavidin HRP (BD Biosciences). After washing, AEC substrate (BD Biosciences) is added before rinsing with water and air-drying. Digitized images are analyzed using CTL ImmunoSpot reader (Cellular Technology Limited). LFn-LACV immunogen specific spots is calculated by subtracting the mean of the negative control values from the mean values of the specific stimulation. Positive responses are greater than four times the mean background, three standard deviations above the background, and >55 spot-forming cells (SFC)/ 2 x 105 total splenic leukocytes.

#### Flow cytometry and ICS assay

Briefly, isolated cells are seeded at a density of 2 x 10□ cells per well in a 96-well U-bottom plate, with LFn-LACV immunogens added to each well at a concentration of 10 μg/ml, followed by incubation at 37°C with 5% CO_2_ for 18 hours. Cells stimulated with PMA will serve as a positive control and cells stimulated with LFn alone as a negative control. Brefeldin A inhibitor is added during the last 4 hours of incubation.

Post-incubation, cells are stained for viability using a viability dye (LIVE/DEAD Fixable Aqua Dead Cell Stain Kit, Thermo Fisher Scientific) and Fc receptors are blocked using anti-mouse CD16/CD32 Fcγ III/II (Thermo fisher sciences) and normal mouse serum. Cells are then stained for extracellular markers using a panel of antibodies (CD8a-Alexa Fluor 488, CD19-Brilliant Violet 650, CD4-APC-eFluor 780, CD3-Alexa Fluor 700 ; all from Thermo Fisher Scientific), washed, and fixed and permeabilized using the FIX & PERM Cell Fixation and Permeabilization Kit (Thermo Fisher Scientific). Following permeabilization, cells are stained for intracellular cytokines (Granzyme B-PE-Cyanine 5.5, IL-2-PE, TNF-alpha-Brilliant Violet 421, IFN-γ-eFluor 660; all from Thermo Fisher Scientific), washed, and resuspended in FACS buffer. Flow cytometry data are acquired using the Aurora system (Cytek® Biosciences) and analyzed using FlowJo software (version 10.2; TreeStar, Inc), retaining live cells and excluding doublets using SSC-A and FSC-A gating.

#### In vivo cytotoxicity assay

Wild-type (C57Bl/6) mice (recipients) were either infected with 10^3^ LACV, or immunized with LACV-Gc or N. Splenocytes (targets) were harvested from donor naïve wild-type mice. RBC were lysed, and the target cells were pulsed with 10 μg/ml of the irrelevant LFn-immunogen (ZIKV), LFn-Gc, LFn-N for 18 h at 37°C. The cells were then washed and labeled with Carboxyfluorescein succinimidyl ester (CFSE) (Invitrogen) in PBS for 10 min at 37°C. LFn-LACV-pulsed cells were labeled with 5 μM CFSE (CFSE^High^) and the irrelevant protein-pulsed cells with 0.5 μM CFSE (CFSE^Low^). After washing, the two cell populations were mixed at a 1:1 ratio and 5 × 106 cells were injected i.v. into naive, infected recipient mice at 14 dpi or immunized weanling mice. After 4 h, the mice were sacrificed and splenocytes were analyzed by flow cytometry, gating on lymphocyte cells using SSC-A and FSC-A gating. The percentage killing was calculated as follows: 100 − ((percentage LACV peptide-pulsed in infected mice/percentage irrelevant peptide-pulsed in infected mice)/(percentage LACV peptide-pulsed in naive mice/percentage irrelevant peptide-pulsed in naive mice)) × 100.

#### LFn-LACV immunizations

1 week old mice were immunized s.c. with 50 μg of each LFn-Gc or LFn-N protein. Mock-immunized mice received LFn-ZIKV-NS3 protein. Mice were boosted with the same dose after 7 days following the first immunization. Mice were then challenged 7 days after second dose immunization with 1000 PFU of LACV. Mice were monitored for neurological signs development.

### QUANTIFICATION AND STATISTICAL ANALYSIS

All statistical analyses were performed using Prism software Version 7.01 (GraphPad) and are described in the figure legends.

### ADDITIONAL RESOURCES

N/A

## REFERENCES

1. Alatrash, R., and Herrera, B.B. (2024). The Adaptive Immune Response against Bunyavirales. Viruses 16, 483.

2. Leventhal, S.S., Wilson, D., Feldmann, H., and Hawman, D.W. (2021). A Look into Bunyavirales Genomes: Functions of Non-Structural (NS) Proteins. Viruses 13, 314.

3. Hulswit, R.J.G., Paesen, G.C., Bowden, T.A., and Shi, X. (2021). Recent Advances in Bunyavirus Glycoprotein Research: Precursor Processing, Receptor Binding and Structure. Viruses 13, 353.

4. Ferron, F., Weber, F., de la Torre, J.C., and Reguera, J. (2017). Transcription and replication mechanisms of Bunyaviridae and Arenaviridae L proteins. Virus Res 234, 118–134. 10.1016/j.virusres.2017.01.018.

5. Westby, K.M., Fritzen, C., Paulsen, D., Poindexter, S., and Moncayo, A.C. (2015). La Crosse Encephalitis Virus Infection in Field-Collected Aedes albopictus, Aedes japonicus, and Aedes triseriatus in Tennessee. J Am Mosq Control Assoc 31, 233–241. 10.2987/moco-31-03-233-241.1.

6. Bennett, R.S., Cress, C.M., Ward, J.M., Firestone, C.Y., Murphy, B.R., and Whitehead, S.S. (2008). La Crosse virus infectivity, pathogenesis, and immunogenicity in mice and monkeys. Virol J 5, 25. 10.1186/1743-422x-5-25.

7. Schneider, C.A., Leung, J.M., Valenzuela-Leon, P.C., Golviznina, N.A., Toso, E.A., Bosnakovski, D., Kyba, M., Calvo, E., and Peterson, K.E. (2024). Skin muscle is the initial site of viral replication for arboviral bunyavirus infection. Nat Commun 15, 1121. 10.1038/s41467-024-45304-0.

8. Haddow, A.D., and Odoi, A. (2009). The incidence risk, clustering, and clinical presentation of La Crosse virus infections in the eastern United States, 2003-2007. PLoS One 4, e6145. 10.1371/journal.pone.0006145.

9. Balfour, H.H., Jr., Siem, R.A., Bauer, H., and Quie, P.G. (1973). California arbovirus (La Crosse) infections. I. Clinical and laboratory findings in 66 children with meningoencephalitis. Pediatrics 52, 680–691.

10. Hartman, A.L., and Myler, P.J. (2023). Bunyavirales: Scientific Gaps and Prototype Pathogens for a Large and Diverse Group of Zoonotic Viruses. The Journal of Infectious Diseases 228, S376–S389. 10.1093/infdis/jiac338.

11. López, K., Wilson, S.N., Coutermash-Ott, S., Tanelus, M., Stone, W.B., Porier, D.L., Auguste, D.I., Muller, J.A., Allicock, O.M., Paulson, S.L., et al. (2021). Novel murine models for studying Cache Valley virus pathogenesis and in utero transmission. Emerg Microbes Infect 10, 1649–1659. 10.1080/22221751.2021.1965497.

12. Winkler, C.W., Myers, L.M., Woods, T.A., Carmody, A.B., Taylor, K.G., and Peterson, K.E. (2017). Lymphocytes have a role in protection, but not in pathogenesis, during La Crosse Virus infection in mice. J Neuroinflammation 14, 62. 10.1186/s12974-017-0836-3.

13. Taylor, K.G., Woods, T.A., Winkler, C.W., Carmody, A.B., and Peterson, K.E. (2014). Age-dependent myeloid dendritic cell responses mediate resistance to la crosse virus-induced neurological disease. J Virol 88, 11070–11079. 10.1128/jvi.01866-14.

14. Basu, R., Nair, V., Winkler, C.W., Woods, T.A., Fraser, I.D.C., and Peterson, K.E. (2021). Age influences susceptibility of brain capillary endothelial cells to La Crosse virus infection and cell death. Journal of Neuroinflammation 18, 125. 10.1186/s12974-021-02173-4.

15. Hefti, H.P., Frese, M., Landis, H., Di Paolo, C., Aguzzi, A., Haller, O., and Pavlovic, J. (1999). Human MxA protein protects mice lacking a functional alpha/beta interferon system against La crosse virus and other lethal viral infections. J Virol 73, 6984–6991. 10.1128/jvi.73.8.6984-6991.1999.

16. Taylor, K.G., and Peterson, K.E. (2014). Innate immune response to La Crosse virus infection. Journal of NeuroVirology 20, 150–156. 10.1007/s13365-013-0186-6.

17. Blakqori, G., Delhaye, S., Habjan, M., Blair, C.D., Sánchez-Vargas, I., Olson, K.E., Attarzadeh-Yazdi, G., Fragkoudis, R., Kohl, A., Kalinke, U., et al. (2007). La Crosse bunyavirus nonstructural protein NSs serves to suppress the type I interferon system of mammalian hosts. J Virol 81, 4991–4999. 10.1128/jvi.01933-06.

18. Pavlovic, J., Schultz, J., Hefti, H.P., Schuh, T., and Mölling, K. (2000). DNA vaccination against La Crosse virus. Intervirology 43, 312–321. 10.1159/000053999.

19. Mukherjee, P., Woods, T.A., Moore, R.A., and Peterson, K.E. (2013). Activation of the innate signaling molecule MAVS by bunyavirus infection upregulates the adaptor protein SARM1, leading to neuronal death. Immunity 38, 705–716. 10.1016/j.immuni.2013.02.013.

20. Winkler, C.W., Myers, L.M., Woods, T.A., Carmody, A.B., Taylor, K.G., and Peterson, K.E. (2017). Lymphocytes have a role in protection, but not in pathogenesis, during La Crosse Virus infection in mice. Journal of Neuroinflammation 14, 62. 10.1186/s12974-017-0836-3.

21. Alatrash, R., Vaidya, V., and Herrera, B.B. (2024). Age-specific dynamics of neutralizing antibodies, cytokines, and chemokines in response to La Crosse virus infection in mice. J Virol 98, e0176224. 10.1128/jvi.01762-24.

22. Lumkong, L., Alatrash, R., Sridhar, S., and Herrera, B.B. (2025). Development of an RT-RPA assay for La Crosse virus detection provides insights into age-dependent neuroinvasion in mice. bioRxiv, 2025.2001.2021.634131. 10.1101/2025.01.21.634131.

23. Herrera, B.B., Tsai, W.Y., Chang, C.A., Hamel, D.J., Wang, W.K., Lu, Y., Mboup, S., and Kanki, P.J. (2018). Sustained Specific and Cross-Reactive T Cell Responses to Zika and Dengue Virus NS3 in West Africa. J Virol 92. 10.1128/jvi.01992-17.

24. Herrera, B.B., Tsai, W.Y., Brites, C., Luz, E., Pedroso, C., Drexler, J.F., Wang, W.K., and Kanki, P.J. (2018). T Cell Responses to Nonstructural Protein 3 Distinguish Infections by Dengue and Zika Viruses. mBio 9. 10.1128/mBio.00755-18.

25. Herrera, B.B., Hamel, D.J., Oshun, P., Akinsola, R., Akanmu, A.S., Chang, C.A., Eromon, P., Folarin, O., Adeyemi, K.T., Happi, C.T., et al. (2018). A modified anthrax toxin-based enzyme-linked immunospot assay reveals robust T cell responses in symptomatic and asymptomatic Ebola virus exposed individuals. PLoS Negl Trop Dis 12, e0006530. 10.1371/journal.pntd.0006530.

26. Akanmu, S., Herrera, B.B., Chaplin, B., Ogunsola, S., Osibogun, A., Onawoga, F., John-Olabode, S., Akase, I.E., Nwosu, A., Hamel, D.J., et al. (2023). High SARS-CoV-2 seroprevalence in Lagos, Nigeria with robust antibody and cellular immune responses. J Clin Virol Plus 3, 100156. 10.1016/j.jcvp.2023.100156.

27. Dakic, A., Shao, Q.-x., D’Amico, A., O’Keeffe, M., Chen, W.-f., Shortman, K., and Wu, L. (2004). Development of the Dendritic Cell System during Mouse Ontogeny1. The Journal of Immunology 172, 1018–1027. 10.4049/jimmunol.172.2.1018.

28. Nelluri, K.D.D., Ammulu, M.A., Durga, M.L., Sravani, M., Kumar, V.P., and Poda, S. (2022). In silico multi-epitope Bunyumwera virus vaccine to target virus nucleocapsid N protein. Journal of Genetic Engineering and Biotechnology 20, 89. 10.1186/s43141-022-00355-y.

29. Shahab, M., Aiman, S., Alshammari, A., Alasmari, A.F., Alharbi, M., Khan, A., Wei, D.-Q., and Zheng, G. (2023). Immunoinformatics-based potential multi-peptide vaccine designing against Jamestown Canyon Virus (JCV) capable of eliciting cellular and humoral immune responses. International Journal of Biological Macromolecules 253, 126678. 10.1016/j.ijbiomac.2023.126678.

30. Adhikari, U.K., Tayebi, M., and Rahman, M.M. (2018). Immunoinformatics Approach for Epitope-Based Peptide Vaccine Design and Active Site Prediction against Polyprotein of Emerging Oropouche Virus. Journal of Immunology Research 2018, 6718083. 10.1155/2018/6718083.

31. Adhikari, U.K., and Rahman, M.M. (2017). Overlapping CD8+ and CD4+ T-cell epitopes identification for the progression of epitope-based peptide vaccine from nucleocapsid and glycoprotein of emerging Rift Valley fever virus using immunoinformatics approach. Infection, Genetics and Evolution 56, 75–91. 10.1016/j.meegid.2017.10.022.

32. Suleman, M., Asad, U., Arshad, S., Rahman, A.u., Akbar, F., Khan, H., Hussain, Z., Ali, S.S., Mohammad, A., Khan, A., et al. (2022). Screening of immune epitope in the proteome of the Dabie bandavirus, SFTS, to design a protein-specific and proteome-wide vaccine for immune response instigation using an immunoinformatics approaches. Computers in Biology and Medicine 148, 105893. 10.1016/j.compbiomed.2022.105893.

33. Maeda, K., West, K., Toyosaki-Maeda, T., Rothman, A.L., Ennis, F.A., and Terajima, M. (2004). Identification and analysis for cross-reactivity among hantaviruses of H-2b-restricted cytotoxic T-lymphocyte epitopes in Sin Nombre virus nucleocapsid protein. Journal of General Virology 85, 1909–1919. 10.1099/vir.0.79945-0.

34. Wang, M., Zhu, Y., Wang, J., Lv, T., and Jin, B. (2011). Identification of Three Novel CTL Epitopes within Nucleocapsid Protein of Hantaan Virus. Viral Immunology 24, 449–454. 10.1089/vim.2011.0026.

35. Wang, M., Wang, J., Kang, Z., Zhao, Q., Wang, X., and Hui, L. (2015). Kinetics and Immunodominance of Virus-Specific T Cell Responses During Hantaan Virus Infection. Viral Immunology 28, 265–271. 10.1089/vim.2014.0135.

36. Schuh, T., Schultz, J., Moelling, K., and Pavlovic, J. (1999). DNA-Based Vaccine against La Crosse Virus: Protective Immune Response Mediated by Neutralizing Antibodies and CD4+ T Cells. Human Gene Therapy 10, 1649–1658. 10.1089/10430349950017653.

37. Boutzoukas, A.E., Freedman, D.A., Koterba, C., Hunt, G.W., Mack, K., Cass, J., Yildiz, V.O., de Los Reyes, E., Twanow, J., Chung, M.G., and Ouellette, C.P. (2023). La Crosse Virus Neuroinvasive Disease in Children: A Contemporary Analysis of Clinical/Neurobehavioral Outcomes and Predictors of Disease Severity. Clin Infect Dis 76, e1114–e1122. 10.1093/cid/ciac403.

38. Case, K.L., West, R.M., and Smith, M.J. (1993). Histocompatibility antigens and La Crosse encephalitis. J Infect Dis 168, 358–360. 10.1093/infdis/168.2.358.

39. Lundegaard, C., Lamberth, K., Harndahl, M., Buus, S., Lund, O., and Nielsen, M. (2008). NetMHC-3.0: accurate web accessible predictions of human, mouse and monkey MHC class I affinities for peptides of length 8-11. Nucleic Acids Res 36, W509–512. 10.1093/nar/gkn202.

40. Calis, J.J., Maybeno, M., Greenbaum, J.A., Weiskopf, D., De Silva, A.D., Sette, A., Keşmir, C., and Peters, B. (2013). Properties of MHC class I presented peptides that enhance immunogenicity. PLoS Comput Biol 9, e1003266. 10.1371/journal.pcbi.1003266.

