## Supplementary Figures for "Immunodominant structural proteins Gc and N drive T cell-mediated protection against La Crosse virus"

**Document S1. Figures S1–S9.**

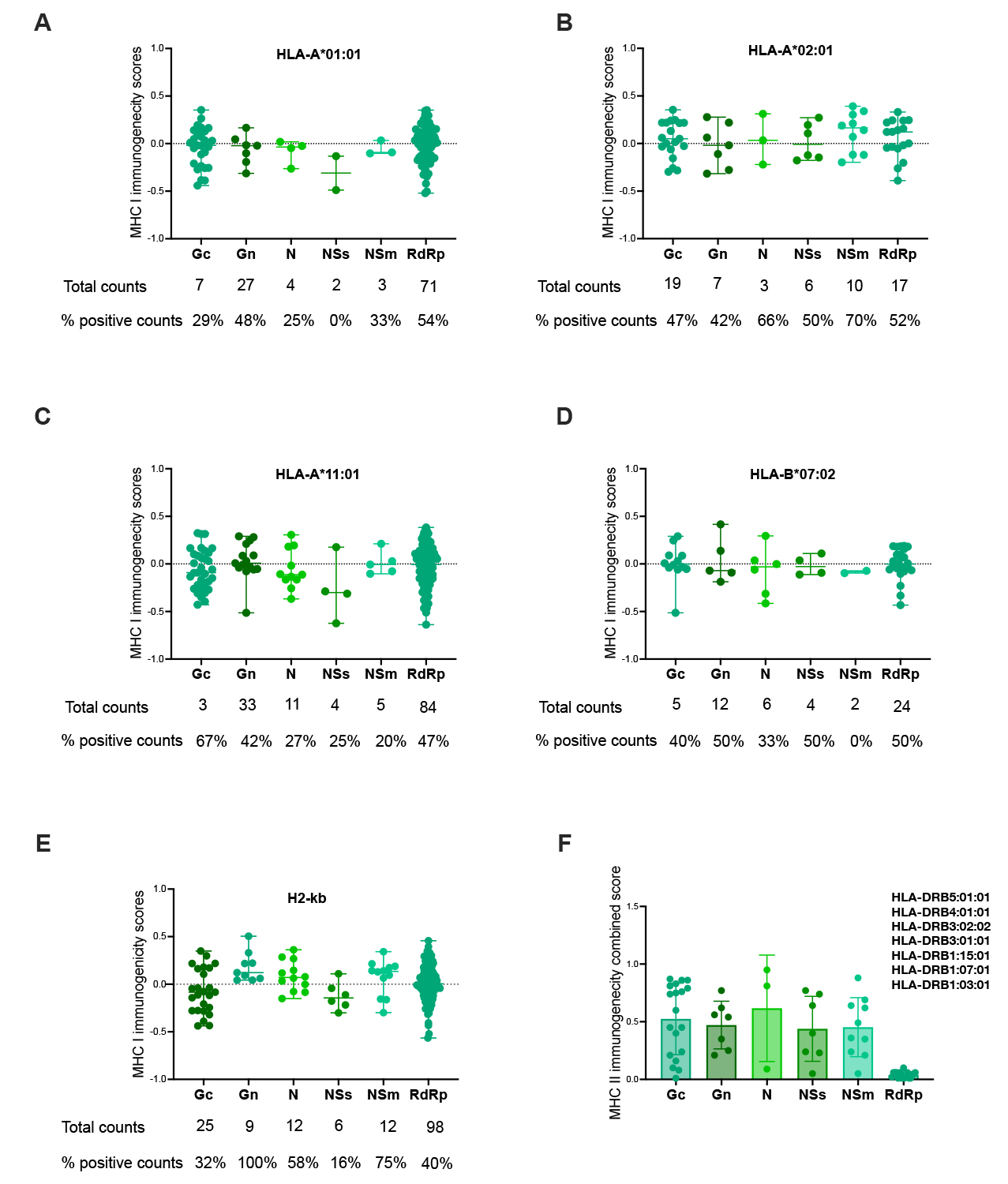

**Figure S1. In silico prediction of immunogenic peptides within LACV proteome with HLA-restriction.** MHC I immunogenicity score of top 1% binders to (A) HLA A*0101, (B) HLA A*0201, (C) HLA A*1101 and (D) HLA B*0702. Number of top 1% of binders and percent immunogenic peptides within each LACV protein is indicated as total counts and counts with positive value of MHC I immunogenicity score respectively. (F) MHC II immunogenicity score of peptides predicted to bind using 7-allele method. Data is presented as individual points with mean ± standard deviation for (A-F).

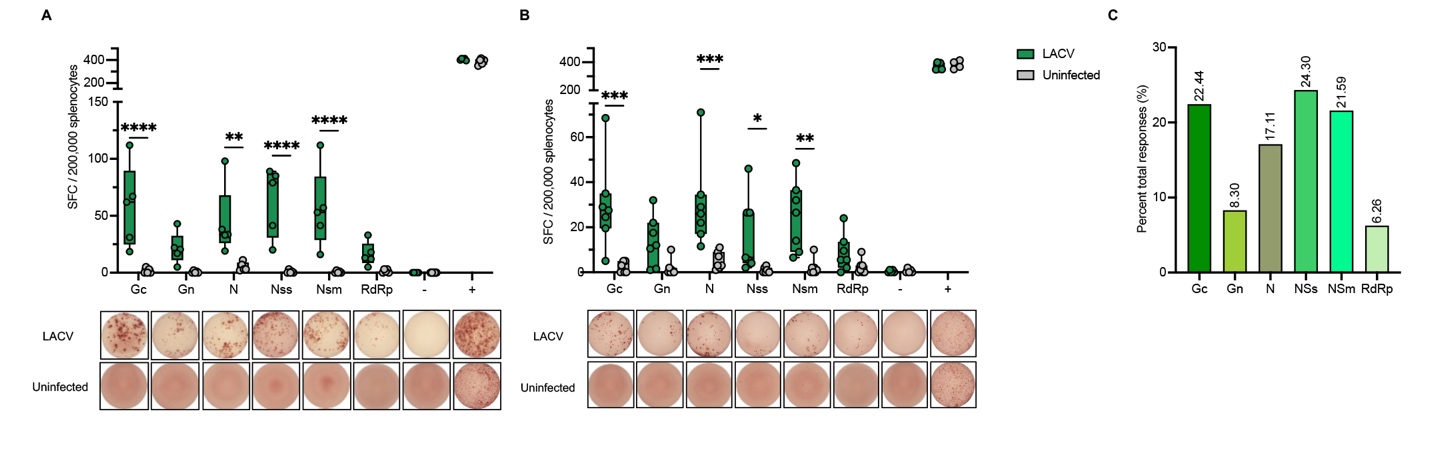
 **Figure S2. LACV-specific ex vivo cellular response in adult wildtype mice.** Top: Quantified spot forming cells (SFC) detected by ELISPOT assay using splenic leukocytes from adult mice infected with LACV at 6 dpi (A) and 14 dpi (B) compared to age-matched non-infected controls. Bottom: Splenic leukocytes were treated with LFn-LACV proteins and IFN-γ responses were detected by ELISPOT assay. (-) negative control: cells stimulated with LFn alone, (+) positive control: cells stimulated with PMA (phorbol 12-myristate 13-acetate). Statistical significance was determined using ANOVA test, *p < 0.05, **p < 0.01, ***p < 0.001, ****p < 0.0001. Data is presented as individual points with minimum and maximum values, (n=5 for A, n=6 for B). (C) Percent of total responses generated against each LFn-LACV protein in adult mice at 6 dpi.

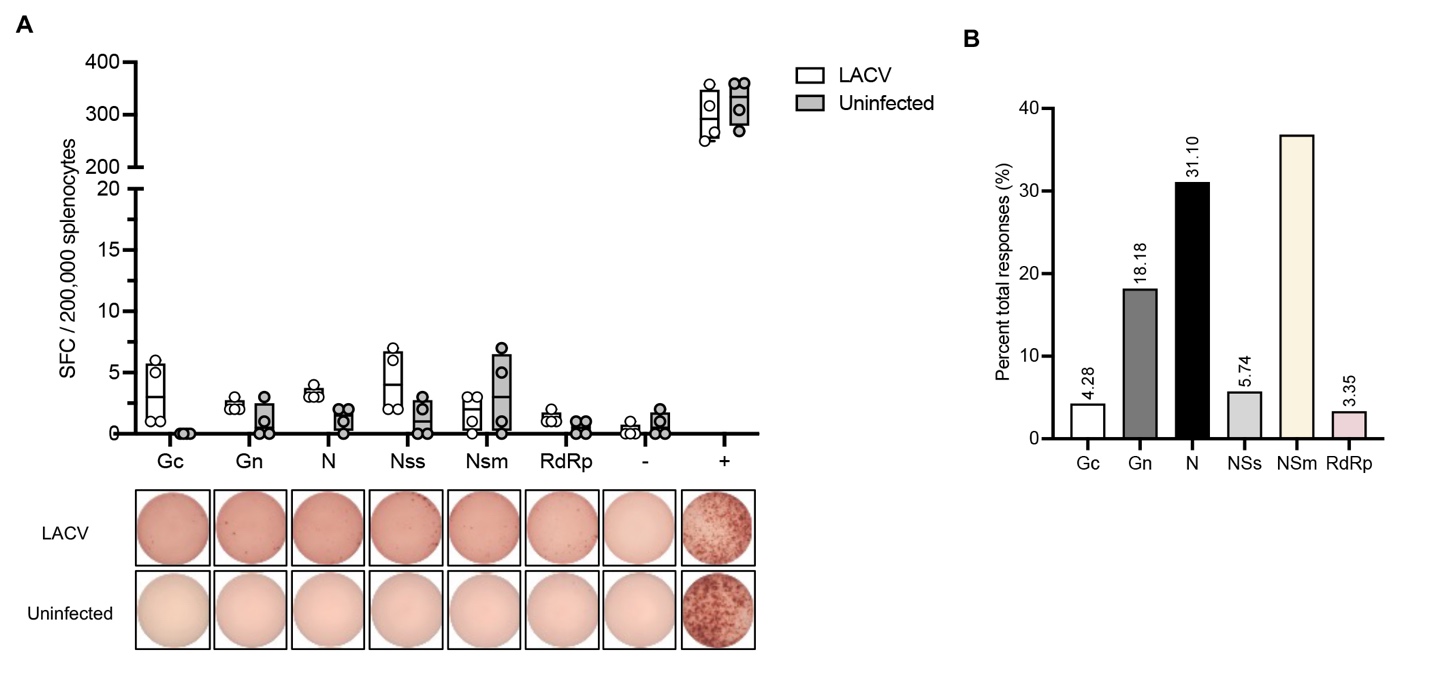

**Figure S3. LACV-specific ex vivo cellular response in weanling wildtype mice.** Top: Quantified spot forming cells (SFC) detected by ELISPOT assay using splenic leukocytes from weanling mice infected with LACV at 6 dpi compared to age-matched non-infected controls. Bottom: Splenic leukocytes were treated with LFn-LACV proteins and IFN-γ responses were detected by ELISPOT assay. (-) negative control: cells stimulated with LFn alone, (+) positive control: cells stimulated with PMA (phorbol 12-myristate 13-acetate). Statistical significance was determined using ANOVA test, *p < 0.05, **p < 0.01, ***p < 0.001, ****p < 0.0001. Data is presented as individual points with minimum and maximum values, (n=5 weanlings, n=4 uninfected). (B) Percent of total responses generated against each LFn-LACV protein in weanling mice at 6 dpi.

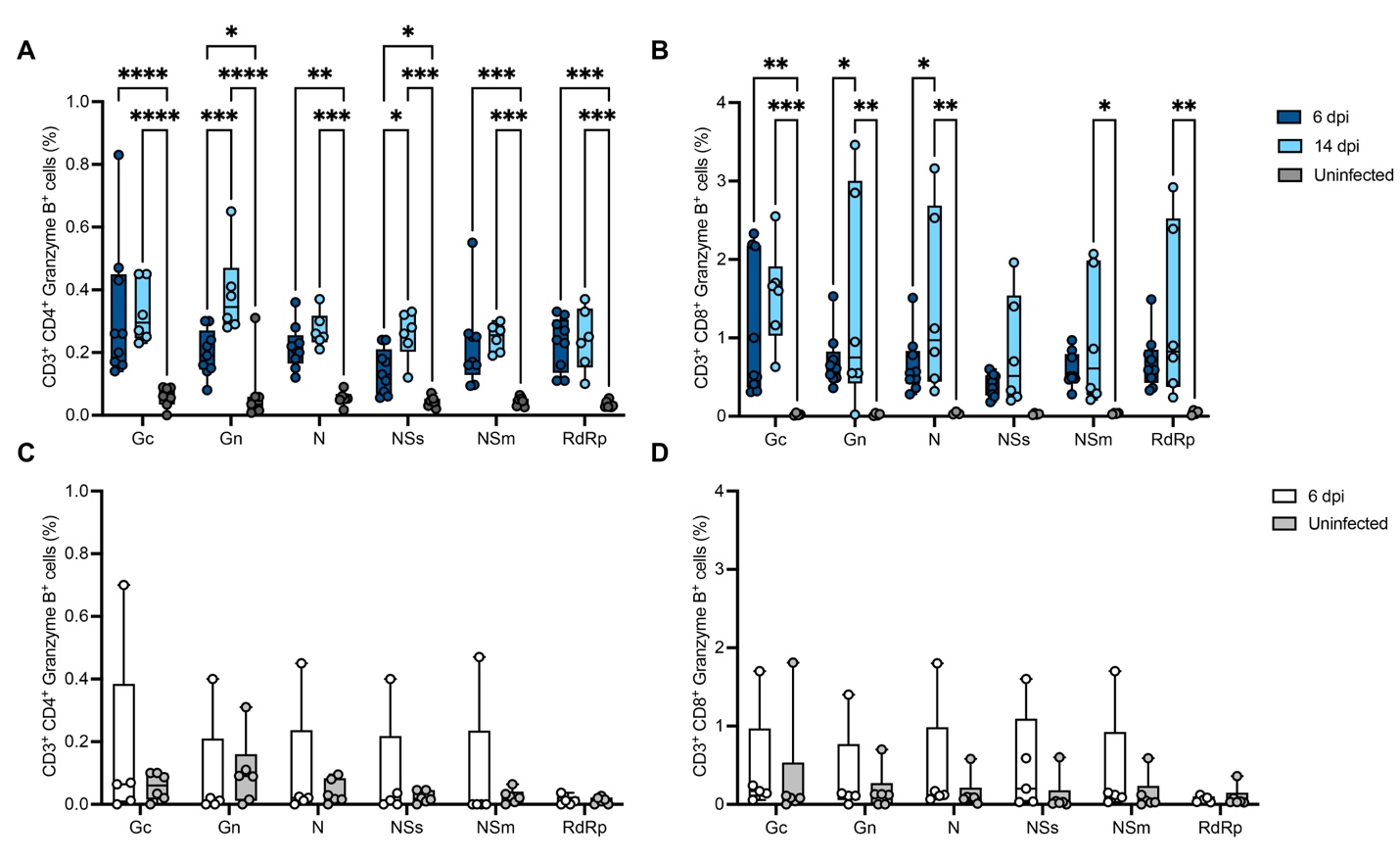

**Figure S4. LACV-specific ex vivo splenic CD4^+^ and CD8^+^ T cell Functional response in LACV infected mice.** Functional LACV-specific CD4^+^ and CD8^+^ T cell responses shown as the percent of CD4^+^ or CD8^+^ cells positive for Granzyme B in CD3^+^ gate in FC/ICS in adult (A, B) and weanling mice (C, D) at 6, 14 dpi compared to age-matched uninfected controls. Data is presented as individual points with minimum and maximum values, (n=9 adults, 5 weanlings).

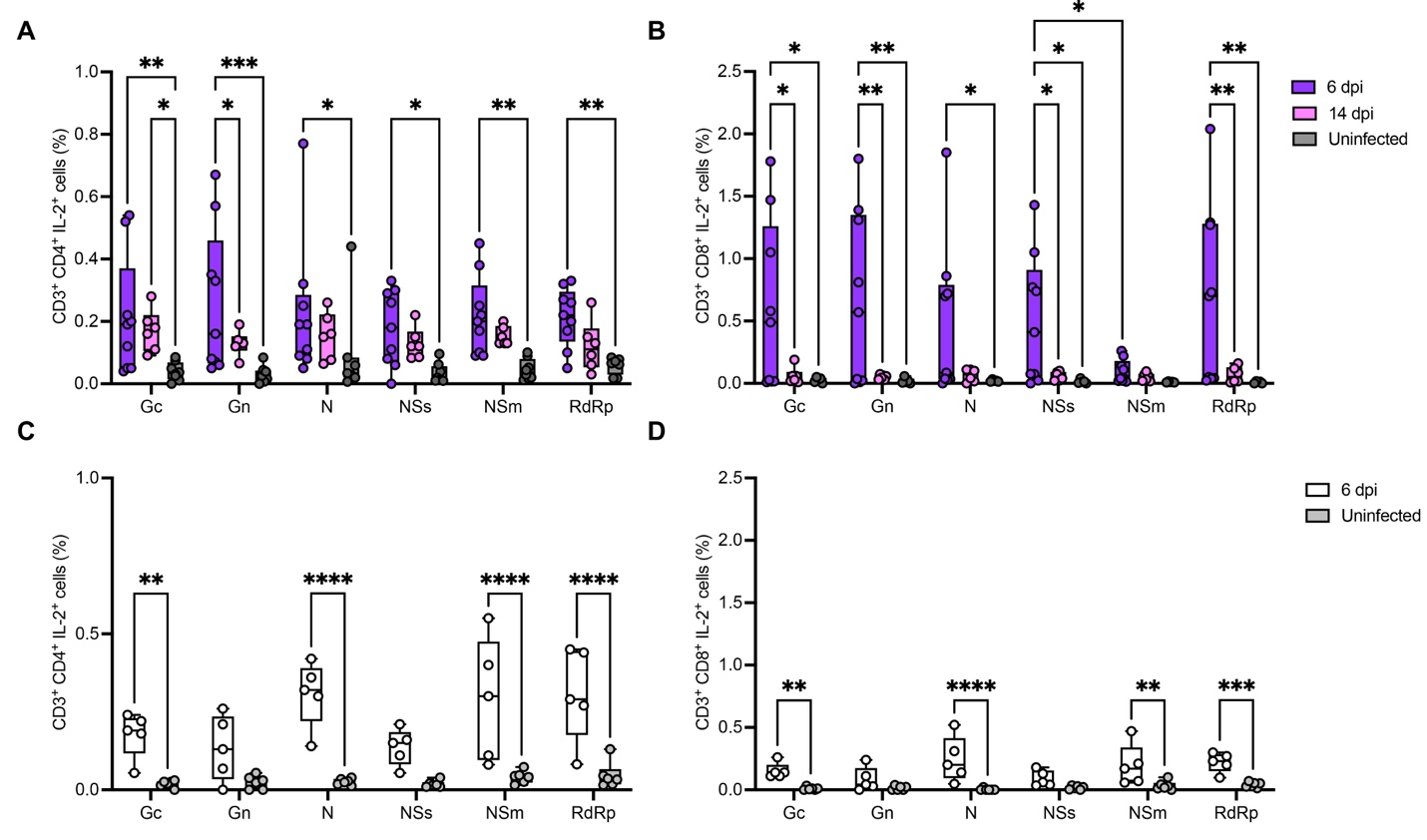

**Figure S5. LACV-specific ex vivo splenic CD4^+^ and CD8^+^ T cell Functional response in LACV infected mice.** Functional LACV-specific CD4^+^ and CD8^+^ T cell responses shown as the percent of CD4^+^ or CD8^+^ cells positive for IL-2 in CD3^+^ gate in FC/ICS in adult (A, B) and weanling mice (C, D) at 6, 14 dpi compared to age-matched uninfected controls. Data is presented as individual points with minimum and maximum values, (n=9 adults, 5 weanlings).

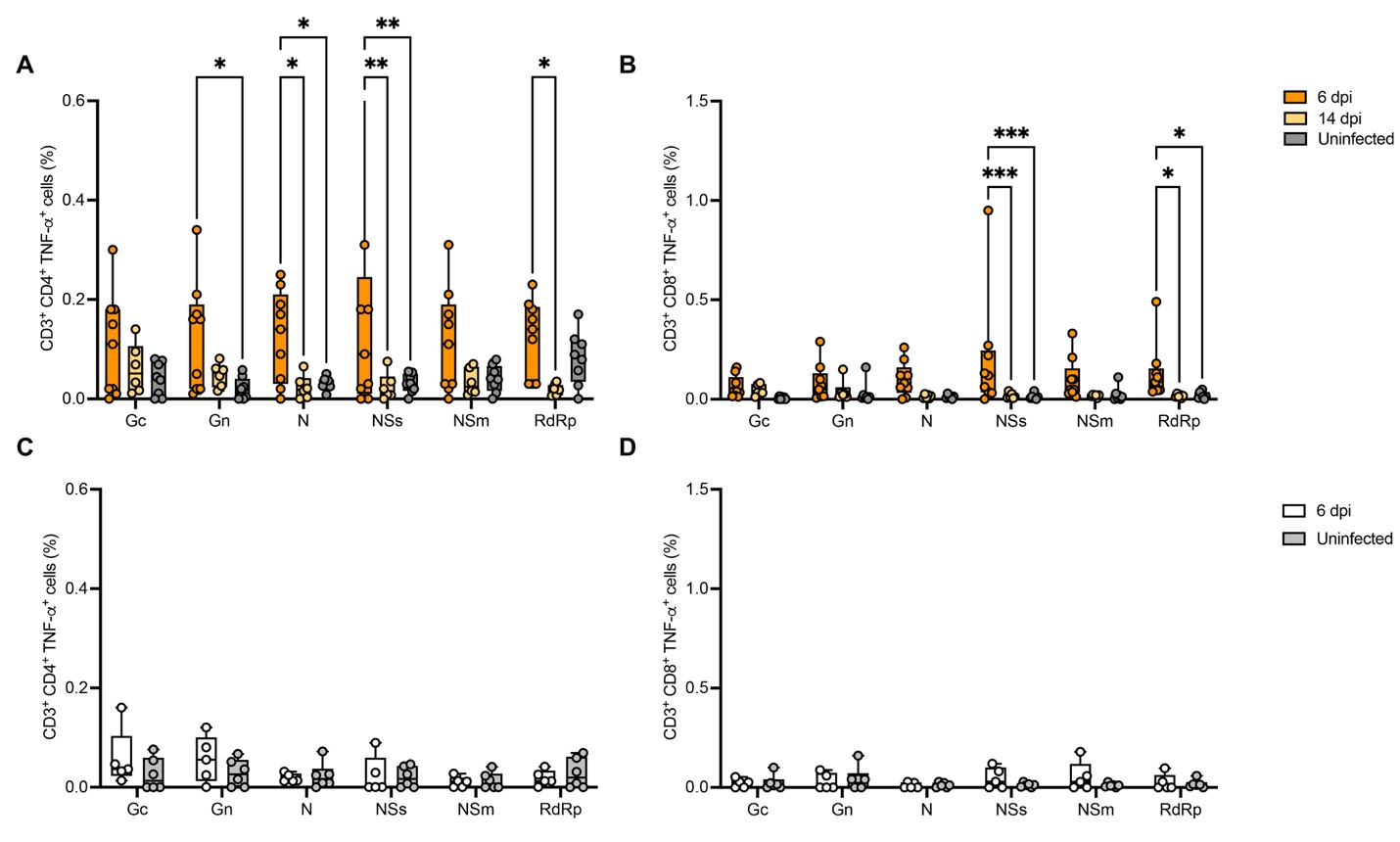

**Figure S6. LACV-specific ex vivo splenic CD4^+^ and CD8^+^ T cell Functional response in LACV infected mice.** Functional LACV-specific CD4^+^ and CD8^+^ T cell responses shown as the percent of CD4^+^ or CD8^+^ cells positive for TNF-ɑ in CD3^+^ gate in FC/ICS in adult (A, B) and weanling mice (C, D) at 6, 14 dpi compared to age-matched uninfected controls. Data is presented as individual points with minimum and maximum values, (n=9 adults, 5 weanlings).

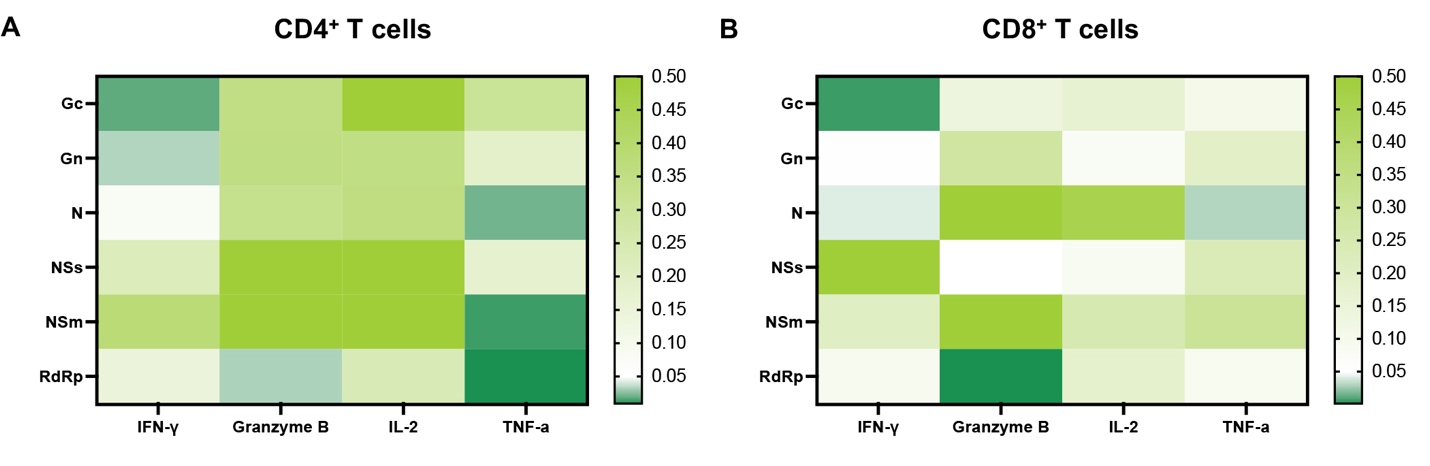

**Figure S7. Functional differences in virus-specific CD4⁺ and CD8⁺ T cells between adult and weanling mice at 6 dpi.** Heatmap illustrating p-values derived from the Mann-Whitney U test comparing the percent of virus-specific CD4⁺ (A) and CD8⁺ T (B) cells producing individual cytokines analyzed in FC/ICS assay in adult versus weanling mice. Columns correspond to the functional markers analyzed, and rows indicate T cell specificity. Darker colors indicate lower p value (n=9 adults, 5 weanlings).

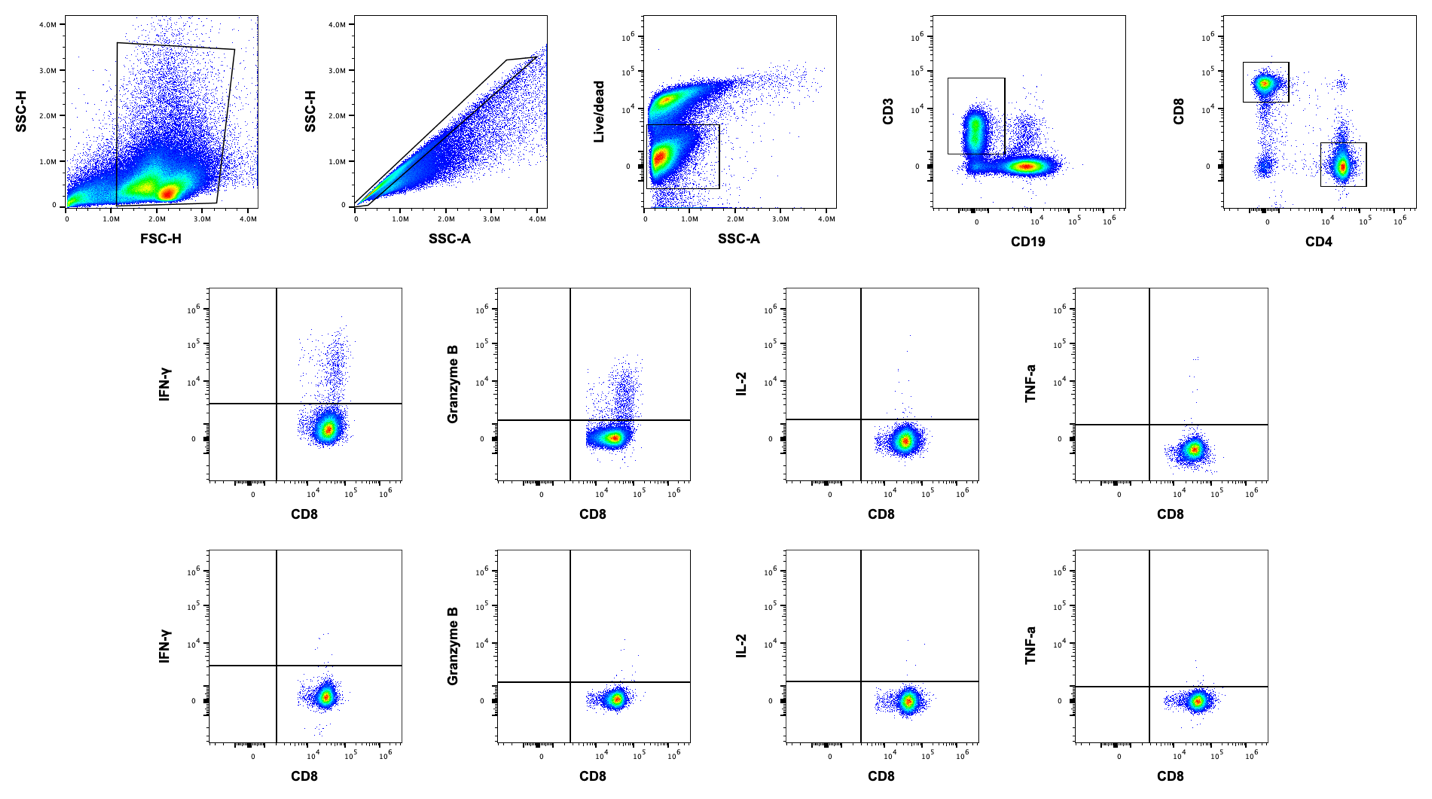
**Figure S8.** Representation of gating strategy for intracellular staining quantification of LACV-specific CD8^+^ Functional responses in adult wildtype mice using ICS/Flow cytometry. Gating for cytokine-positive populations is shown as an example for infected and uninfected adult mice at 14 dpi.

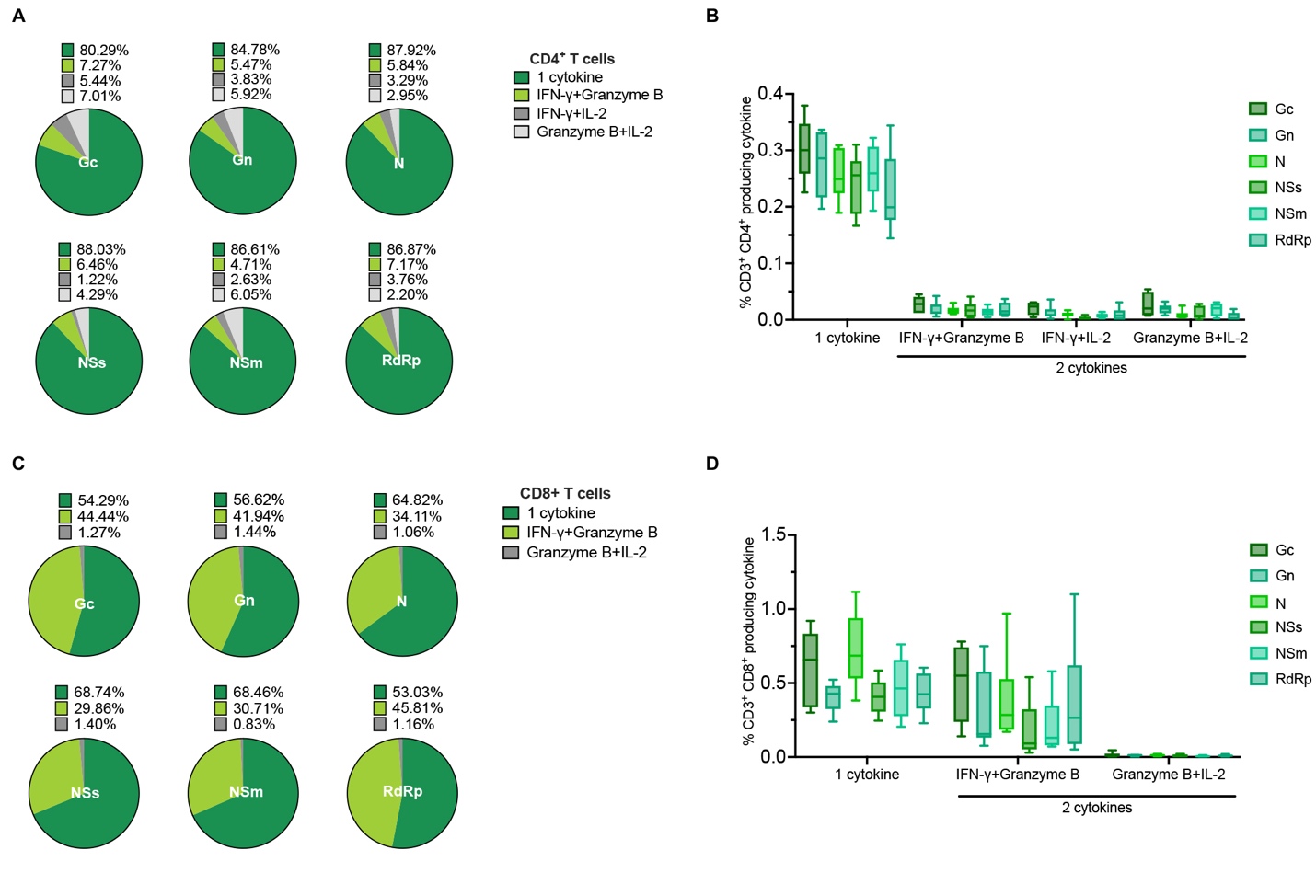

**Figure S9. Polyfunctional profiles of LACV-specific CD4⁺ and CD8⁺ T cells at 14 dpi.** Pie charts depicting the distribution of LACV-specific CD4⁺ T cell (A) and CD8^+^ T cell (C) populations with single or dual functional profiles, defined by cytokine production. (B, D) Bar graph showing the total percentage of LACV-specific CD4⁺ T or CD8^+^ T cells producing one or two cytokines in response to infection. Data is presented as mean with minimum and maximum values (n=9).
